# Validated machine learning algorithm with sub-clonal sensitivity reveals widespread pan-cancer human leukocyte antigen loss of heterozygosity

**DOI:** 10.1101/2021.05.20.445052

**Authors:** Rachel Marty Pyke, Dattatreya Mellacheruvu, Steven Dea, Charles W. Abbott, Simo V. Zhang, Lee McDaniel, Eric Levy, Gabor Bartha, John West, Michael P. Snyder, Richard Chen, Sean Michael Boyle

**Affiliations:** Personalis, Inc., Menlo Park, CA; Stanford University, Palo Alto, CA

## Abstract

Human leukocyte antigen loss of heterozygosity (HLA LOH) allows cancer cells to escape immune recognition by deleting HLA alleles, causing the suppressed presentation of tumor neoantigens that would otherwise bind to them. Despite its importance in immunotherapy response, few methods exist to detect HLA LOH, and their accuracy is not well understood. Here, we develop DASH (**D**eletion of **A**llele-**S**pecific **H**LAs), a novel machine learning-based algorithm to detect HLA LOH from paired tumor-normal sequencing data. Through validation with cell line mixtures and patient-specific digital PCR, we demonstrate increased sensitivity compared to previously published tools and pave the way for clinical utility. Using DASH on 611 patients across 15 tumor types, we found that 18% of patients had HLA LOH. Moreover, we show inflated HLA LOH rates compared to genome-wide LOH and correlations between CD274 (PD-L1) expression and MSI status, suggesting the HLA LOH is a key immune resistance strategy.

## Introduction

The success of immune checkpoint blockade (ICB) therapies confirms that the immune system is capable of recognizing and eliminating cells that present tumor neoantigens via major histocompatibility complexes (MHC) ^1^. However, a large percentage of patients do not respond to these therapies, driving the need for new methods that can predict response ^2^. Elucidating the cause of resistance has proven more challenging than initially anticipated because of the complex tumor intrinsic and extrinsic mechanisms underlying ICB evasion ^3,4^.

Antigen presentation can influence the evolution of tumors ^5–7^. Each individual has up to six different classical class I human leukocyte antigen (HLA) alleles capable of presenting a unique set of antigens to the immune system. Tumor mutations are enriched in HLA and B2M genes, suggesting that the functionality of these genes is critical to mounting an immune response ^8–10^. Germline sequence diversity of HLA alleles also impacts tumor evolution; moreover, the impact of this evolution appears to be even more pronounced in the presence of ICB ^11,12^. Thus, somatic LOH in the HLA region has the potential to cause extreme reduction in HLA sequence diversity and is increasingly being recognized as a pathway of ICB escape ^13^ and an indicator of response ^14^.

Detecting HLA LOH accurately from sequencing data is of interest given the growing ubiquity of tumor molecular profiling. Many copy number detection algorithms estimate tumor purity and ploidy to establish the expected relationships between b-allele frequency (sequencing depth ratio between two alleles - A and B), genomic sequencing depth and copy number deletions ^15,16^. However, general copy number detection algorithms used for detecting HLA LOH from sequencing data have proven unreliable because the polymorphic nature of the genes causes poor alignment to the reference genome. Moreover, the complexity of the sequence variation obscures the specific HLA allele that has been deleted, information crucial for neoantigen therapy design. Though genomic interrogation of the HLA region has been a challenge across applications, both alignment to allele-specific references and application of genome graphs have improved HLA typing and HLA somatic mutation identification ^8,17,18^. Although allele-specific alignment has been used for several HLA LOH detection algorithms, these algorithms are still based on tumor-only sequencing ^19^ or on standard copy number variant (CNV) algorithms that fail to account for HLA-specific challenges such as differences in exome probe capture between alleles ^15^. Furthermore, CNV detection tools are notoriously poor for samples with low tumor purity and have trouble detecting subclonal deletions, raising concerns regarding the sensitivity and accuracy of these tools ^14^. Thus, despite growing interest in the field, many studies still supplement deletions in regions surrounding the HLA region as a proxy for HLA LOH instead of using an HLA LOH specific algorithm and forfeit information about the identity of the deleted allele ^12,20,21^.

Validating performance of HLA LOH detection algorithms has been an additional hurdle in the field. Two main approaches have been taken. First, concordance has been assessed between HLA LOH calls and copy number calls made by a standard CNV algorithm in regions flanking each HLA gene ^16^. Second, primers have been designed to capture the regions surrounding the HLA genes and PCR has been used to test for copy number loss in patients predicted to have all HLA loci impacted. However, neither of these approaches validate the specific allele lost nor address the accuracy of calls for low tumor purity samples or samples with subclonal HLA LOH ^22^, hindering their utility as orthogonal validation approaches for securing clinical approval.

To more accurately profile HLA allele-specific LOH, we designed a machine learning algorithm, DASH (**D**eletion of **A**llele-**S**pecific **H**LAs), to capture the unique features of the HLA region and demonstrated the limit of detection for sub-clonal events and low tumor purity using sequencing data mixtures from cell lines. Moreover, we developed an allele-specific digital PCR assay to orthogonally validate DASH and probed the mechanistic impact of HLA LOH on peptide presentation with quantitative immunopeptidomics. Using a cohort of 611 patients spanning 15 tumor types, a range of tumor purity and all clinical grades, we applied DASH and found widespread HLA LOH, with the majority of tumor types having at least 20% of patients impacted. Finally, we found evidence for the evolutionary selection of HLA LOH, suggesting its importance in tumor development and immune escape.

## Results

### DASH employs machine learning with novel features to detect HLA LOH detection

Traditional copy number detection tools are unable to accurately detect allele-specific deletions in the HLA region; thus, we developed a machine learning approach to overcome the unique challenges this region presents. To create a dataset for learning genomic features important to accurately calling allele-specific LOH in HLA genes, we profiled 279 paired tumor and healthy normal samples (either adjacent tissue or PBMC) from patients. To type the HLA alleles, we used whole exome sequencing on the ImmunoID NeXT Platform, which has additional capture probes covering specific HLA alleles and has been validated for accurate HLA typing (**Figure 1A, Supp. Table 1**). After determining the HLA types for each patient, we mapped all HLA-specific reads to each patient-specific HLA reference with stringent alignment parameters ^23^. From the alignment data, we calculated allele-specific sequencing depth and developed four features based on the genomic positions containing allele-specific germline variations (**Figure 1B**). First, we wanted to capture a measurement of b-allele frequency (allelic fraction), but we found that the sequencing depth in the data from adjacent normal tissue was often different between alleles. To address this issue, we divided the tumor b-allele frequency by the adjacent normal b-allele frequency, creating an adjusted b-allele frequency (**Supp. Figure 1A**). Second, we calculated an allele-specific sequencing depth ratio by dividing the tumor depth of an allele by the adjacent normal depth of the same allele and normalizing by the depth across the rest of the exome (**Supp. Figure 1B**). Third, we found that alleles with consistently lower sequencing depth than the alternate allele across the entire gene were likely to be deleted, whereas sporadically lower depth may be due to stochastic variation (p<2.2e-14, paired T test with the null hypothesis that samples with and without HLA LOH would have the same consistency of depth across the gene, **Supp. Figure 1C**). Thus, we added a feature, consistency of sequencing depth, to capture this observation. Finally, in order to distinguish allelic imbalance driven by LOH from allelic imbalance driven by a large amplification of the alternate allele, we added a feature, total sequencing depth ratio, that captures the combined depth of the two alleles. We found that patients with HLA LOH have significantly lower total sequencing depth ratios (p=0.0004, paired T test with the null hypothesis that patients with and without HLA LOH will have the same distribution of total depth ratios, **Supp. Figure 1D**).

**Figure 1.**
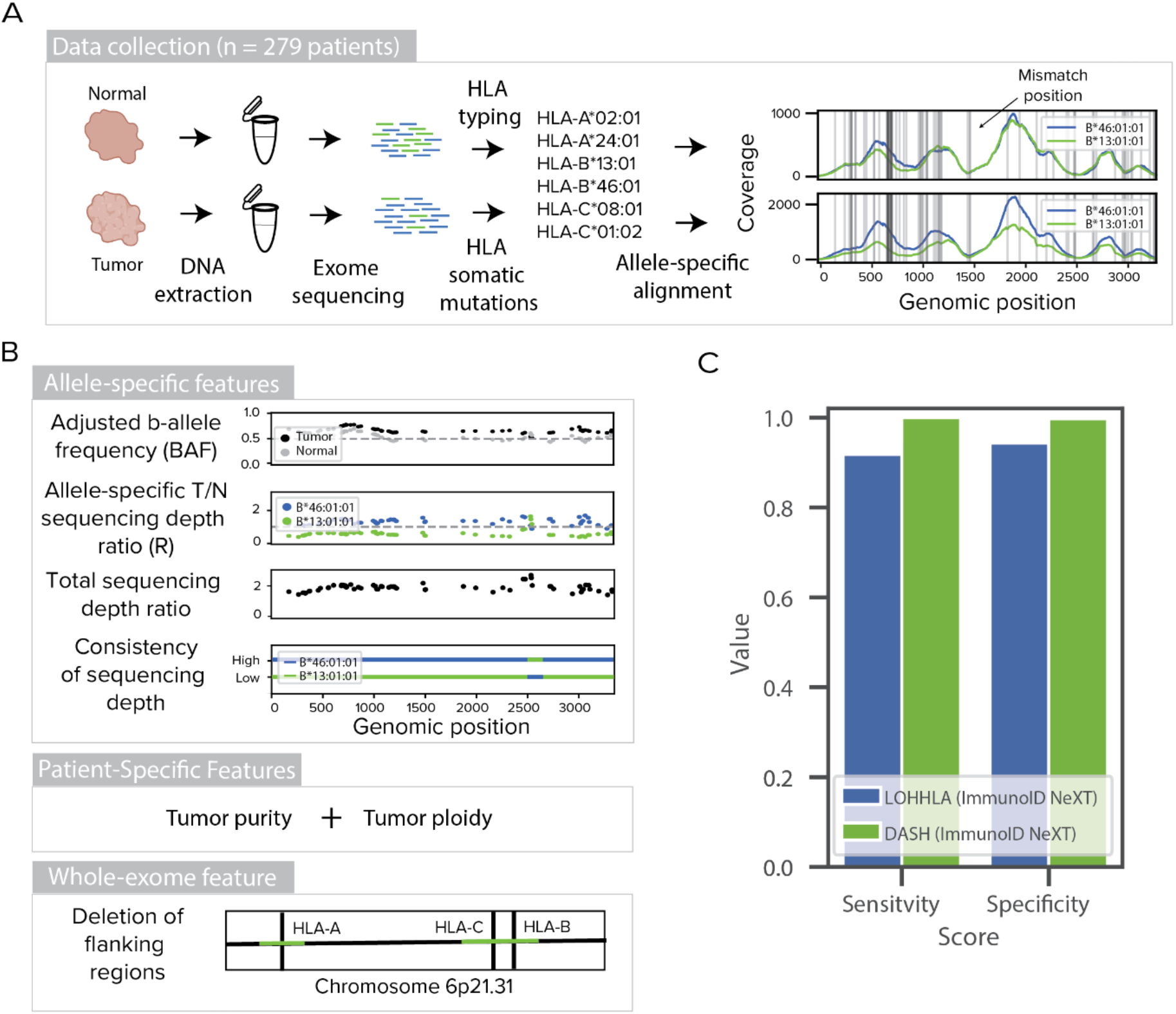
Training, performance and limit of detection of DASH algorithm. (**A**) Tumor and adjacent normal samples were collected for 279 patients. DNA was extracted and exome sequencing was performed. HLA typing was derived from the normal sample and HLA somatic mutations were obtained from the tumor sample. Reads mapping to an HLA reference database were assembled and mapped onto the patient-specific HLA alleles. Horizontal grey lines denote positions where the homologous alleles differ from each other. (**B**) Features used to train the DASH algorithm include the adjusted b-allele frequency, sequencing depth ratio, total sequencing depth ratio, consistency of sequencing depth, tumor purity, tumor ploidy and deletion of flanking regions. Dashed grey lines represent the expected values for a sample without copy number change. All features are used to train an XGBoost prediction model. (**C**) A bar plot showing the sensitivity and specificity of HLA LOH detection by DASH and LOHHLA for samples in the test data set. Both algorithms were tested on the HLA-enhanced ImmunoID NeXT Platform. Only samples with at least 20% tumor purity are included.

To supplement the allele-specific features, we also included two patient-specific features, tumor purity and tumor ploidy ^15^, that are identical across HLA-A, HLA-B and HLA-C of a particular patient (**Supp. Figure 1E-F**). Finally, since 73% of copy number alterations causing HLA LOH are deletions of greater than one megabase (compared to 63% of genome-wide copy number alterations being greater than one megabase, **Supp. Figure 1G**), we reasoned that the genomic regions flanking the genes of interest can provide useful information to supplement the within-gene data. Thus, we utilized the whole exome nature of our dataset to generate a final feature, deletion of flanking regions, that measures deletions in the 10 kb region surrounding each gene.

### DASH outperforms competing algorithms in cross validation evaluation

From the 279 patients, we collected these six features for 720 heterozygous HLA loci, excluding homozygous alleles because LOH cannot occur. To add training labels, we visualized all 720 alleles and manually curated deleted alleles and identified that 19.6% of the heterozygous loci had HLA LOH (**Supp. Figure 2A-B**, see Methods). Though generic LOH calling tools are unable to identify the specific allele that has been lost, we first wanted to assess the ability of a generic LOH calling tool to detect a deletion in HLA genes as a baseline on ImmunoID NeXT. Using Sequenza, we found 92.9% specificity and 95.0% sensitivity (F1-Score = 0.848; **Supp. Figure 3A**). Then, using 10-fold cross validation and the features outlined above, we trained a gradient boosted regression (XGBoost) algorithm, which outperformed several other algorithms, to predict deleted alleles (**Supp. Figure 3B**). DASH reaches 98.7% specificity and 92.9% sensitivity while LOHHLA, another published HLA LOH algorithm, only achieves 94.3% specificity and 78.8% sensitivity (F1-Scores = 0.939 and 0.777 respectively; DASH AUROC=0.939, DASH AUPRC=0.940; **Supp. Figure 3C-E**). Though Sequenza is unable to detect the lost allele, we observe that Sequenza outperforms LOHHLA, but not DASH, in its ability to detect a deletion.

Of note, the majority of the incorrect calls have low tumor purity, highlighting the difficulty of accurately predicting HLA LOH at low tumor purity levels (**Supp. Figure 3F**). When samples with tumor purity below 20% are removed from the analysis, DASH continues to outperform LOHHLA at 99.7% specificity and 100% sensitivity compared to 94.3% specificity and 91.8% sensitivity (F1-Scores=0.995 and 0.857 respectively; DASH AUROC=0.990, DASH AUPRC=0.992; **Figure 1C**). The algorithm revealed that all seven of our features were independently contributing to the model, with deletion of flanking regions and adjusted BAF impacting the outcome most significantly (**Supp. Figure 4A**). We also found that none of DASH’s features alone could achieve this level of performance (**Supp. Figure 4B**).

All three of these algorithms were tested on the HLA-enhanced ImmunoID NeXT Platform, suggesting that performance may decrease with other exome platforms. One of the differences between the ImmunoID NeXT Platform and other exomes for this application is the boosted sequencing of the HLA loci. ImmunoID NeXT provides an exonic region sequencing depth of 1586x prior to stringent postprocessing via DASH and a full gene sequencing depth of 587x after post-processing, while typical cancer exomes are performed at lower sequencing depths. For example, example cohorts from dbGaP provide sequencing depths of 373x and 161x before and after post-processing, respectively (**Supp. Figure 5A-B**) ^24,25^. Thus, we evaluated the performance of DASH with decreasing sequencing depths across the HLA genes. We found that the F1-score drops slightly as sequencing depth decreases, dropping .06 from the ImmunoID NeXT depth to 100x depth (**Supp. Figure 5C**). Furthermore, the performance is highly dependent on tumor purity, with the F1-score dropping more dramatically at lower sequencing depths for samples with lower tumor purity (**Supp. Figure 5D**).

### In silico cell line mixtures demonstrate the sub-clonal sensitivity of DASH

Though more study is required to understand the general timing of HLA LOH, it has been observed in some instances as occurring late in tumor progression as a resistance mechanism ^13,22^. Furthermore, tumor types that tend to be most responsive to ICB (lung, skin) also tend to produce lower purity samples ^26^. Finally, assessment of performance with manually curated labels has some limitations. For these three reasons, we thought that it was critical to assess the limit of detection of DASH with gold standard samples in both sub-clonal and low tumor purity settings. We sequenced 22 tumor-normal paired cell lines with ImmunoID NeXT and identified four with HLA LOH in at least one locus (**Supp. Table 2, Supp. Figures 6-8**). For three of these cell lines (CRL-5911, CRL-5922 and CRL-2314), we sub-sampled the tumor sequencing data and mixed it with complementary normal sequencing data to achieve simulated purity levels in replicates of ten (see Methods, **Figure 2A, Supp. Table 3**).

**Figure 2.**
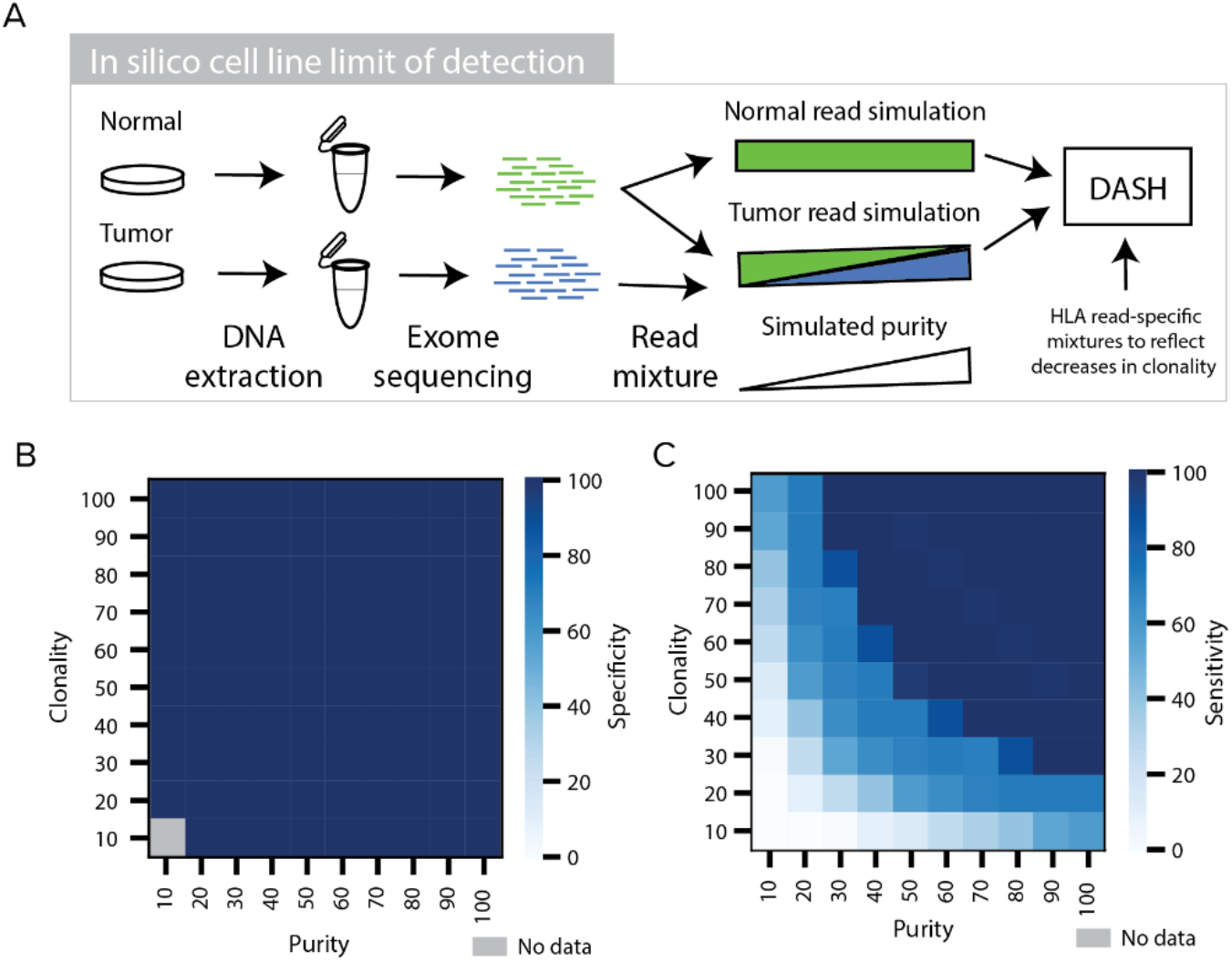
*In silico* cell line mixtures to determine DASH limit of detection. (**A**) A schematic showing the tumor and normal cell line mixing approach for simulating low purity sample pairs. (**B-C**) Heatmaps showing the (**B**) specificity and (**C**) sensitivity of DASH to capture HLA LOH in simulated samples of differing purity and clonality. Dark blue denotes high sensitivity or specificity, light blue denotes low sensitivity or specificity and grey denotes no data.

With all three cell lines analyzed together, greater than 98% sensitivity was observed at all tumor purity levels higher than 27% while greater than 98% specificity was observed across all tumor purity levels (**Supp. Figure 9A-B**). While LOHHLA reached the same level of specificity as DASH, LOHHLA shows weaker sensitivity, with greater than 97% above 35% tumor purity (**Supp. Figure 9C-D**). Notably, performance is much higher for both algorithms in two of the cell lines (CRL-5911 and CRL-2314) than the third (CRL-5922), with 100% sensitivity for tumor purities above 8% tumor purity (**Supp. Figure 10A-D, I-L and Q-T**). Next, we mixed the HLA-mapping reads with the whole exome reads across a range of different ratios to simulate the potential spectrum of tumor purities and sub-clonalities (see Methods). Both DASH and LOHHLA retained greater than 99% specificity across all sub-clonality and tumor purity levels (**Figure 2B, Supp. Figure 9E**). As expected, both their sensitivities decreased with lower purity and clonality; however, DASH still retained greater than 97% sensitivity until the mixture of reads was at least 25% derived from the HLA LOH event (as observed in the tumor purity and HLA LOH clonality combination: 50% purity and 50% clonality). In contrast, LOHHLA was unable to recognize the vast majority of events that were less than 80% clonal, highlighting the accuracy and low detection limit of DASH to capture sub-clonal HLA LOH events (**Figure 2C, Supp. Figure 9F, Supp. Figure 10E-H, M-P, U-X**).

### Genomic validation of DASH in tumor samples using allele-specific digital PCR with patient-specific primer designs

After establishing DASH’s ability to detect HLA LOH in challenging samples, we sought to orthogonally validate our prediction ability with digital PCR (dPCR). Due to the highly polymorphic nature of HLA alleles, PCR primers and probes had to be independently designed for each pair of alleles in each patient (**Figure 3A**, Supplemental Data). In addition to having to target the alleles unique to each patient, the probes were also designed to avoid targets on other patient-specific alleles. As a proof of concept and to determine the limit of detection of the dPCR assay, we designed primers for the predicted retained and lost HLA-C alleles for one of the cell lines previously interrogated with *in silico* mixtures (CRL-5911). Then, we mixed the tumor and blood (normal) cell line DNA to emulate decreasing levels of tumor purity in triplicate and performed dPCR with both allele-specific designs. Importantly, we also multiplexed the allele primers with RNase P (RPP25) primers as a diploid control. To ensure the specificity of the primers, we first confirmed that both the predicted lost and predicted retained allele copies (normalized by half of the diploid RNase P copies) resulted in a single copy in the normal sample with a tumor purity of zero (**Figure 3B, Supp. Figure 11A**). Then, we compared the copy number of each allele in the tumor sample to the normal sample with the null hypothesis that the tumor sample would have greater than or equal copy number than the normal sample. As anticipated, we found zero copies of the lost allele in the tumor sample, confirming the allele-specific LOH event, and dPCR sensitivity for 10% tumor purity and above, confirming the sensitivity and reproducibility of allele-specific dPCR as an orthogonal method (**Figure 3B, Supp. Figure 11A**).

**Figure 3.**
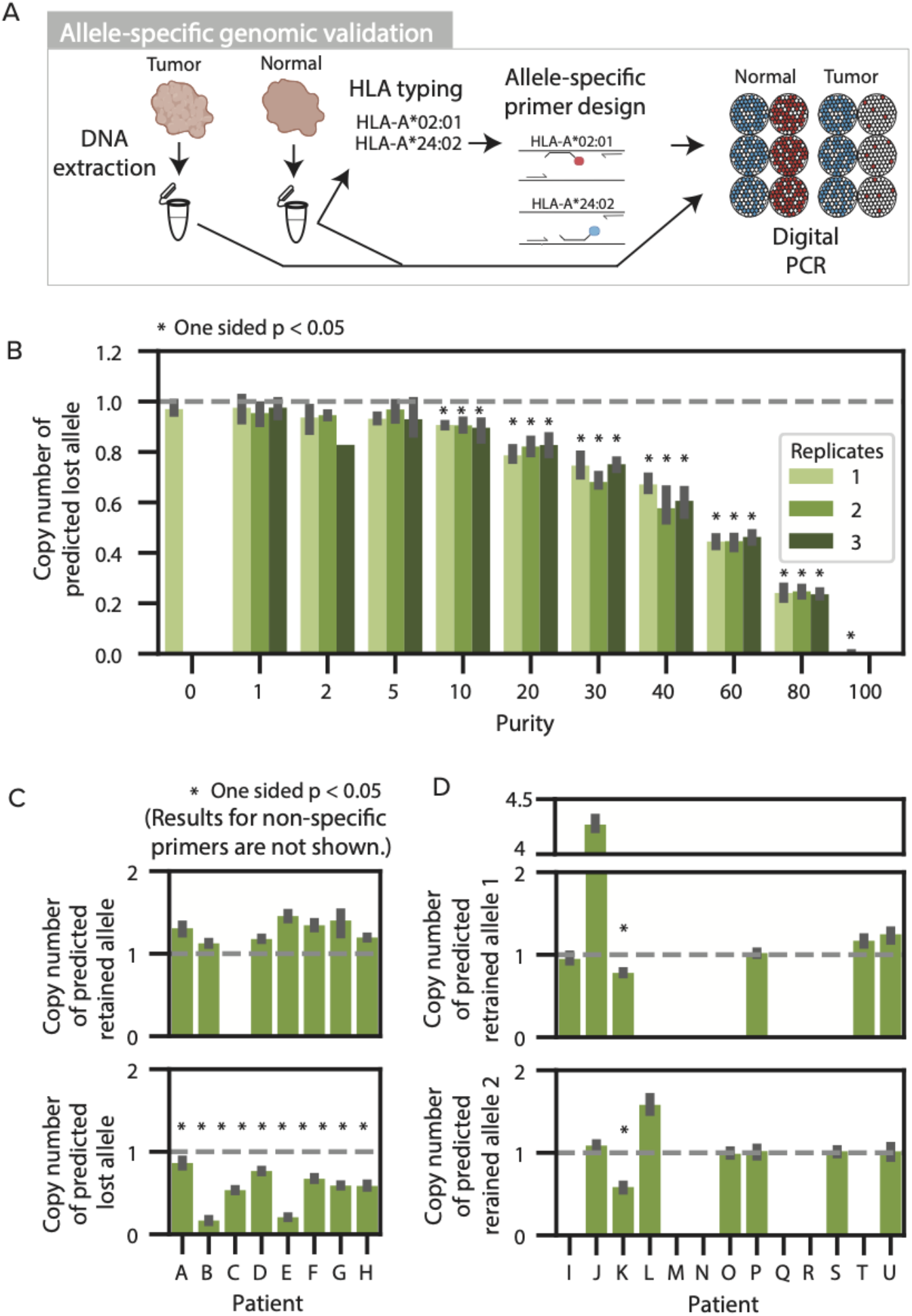
Allele-specific genomic validation with digital PCR. (**A**) Allele-specific genomic validation was performed using paired tumor and adjacent normal fresh frozen samples. DNA was extracted from each sample. Allele-specific primers were designed specifically for each patient. Digital PCR was performed on each sample to orthogonally determine the allele-specific copy number. (**B**) Bar plot showing the allele-specific copy number of the predicted lost allele, relative to RNase P, as measured by dPCR for cell line mixtures of varying tumor purities. The dashed line denotes the expected value for no change in copy number. Asterisks denote a statistically significant difference from the copy number in the normal sample with a one-sided Student T test. (**C**) Bar plots denoting the HLA allele dPCR copy number relative to the multiplexed RNase P for patient tumor samples. The alleles predicted by DASH to be retained are shown on the top plot while the alleles predicted to be deleted are shown on the bottom plot. The dashed grey lines denote the expected copy number of one if there are no copy number alterations. Asterisks denote samples with p-values less than 0.05 as determined by a one-sided Student T test. (**D**) Bar plots showing the HLA allele dPCR copy number relative to the multiplexed RNAse P for patient tumor samples where both alleles were predicted to be retained.

Since the detection of HLA LOH in the cell line with dPCR was successful, we profiled a set of 21 patient tumors: 4 confirmatory samples from the training data and 17 independent samples. Of these 21 samples, 8 were predicted to have HLA LOH by both DASH and LOHHLA, 1 was predicted to have HLA LOH by LOHHLA and not DASH, and 10 were predicted not to have HLA LOH by either algorithm. We designed primers for the predicted retained and lost alleles (**Supp. Table 4**) and performed dPCR on DNA from normal (adjacent) and tumor samples (**Supp. Figure 11B-E, Supp. Table 5**). First, we ensured the specificity of the allele-specific primers (**Supp. Figure 12A-B**). We found that the RNase P-adjusted copy number for 30 of the 44 primers were highly specific and in close proximity to one copy; however, we excluded several alleles due to low specificity. Next, we evaluated the RNase P-adjusted copy number in the tumor samples. For the samples with predicted HLA LOH, we found a significant reduction of copy number in all eight of the predicted lost alleles by DASH and no reduction in copy number in all of the predicted retained alleles with specific primers in the tumor samples (**Figure 3C**). For the samples predicted to not have HLA LOH by both algorithms, none of the alleles showed a significant reduction in copy number by digital PCR (**Figure 3D**). Finally, for the sample predicted to have HLA LOH by LOHHLA but not by DASH (Patient K), both alleles appear to have a reduction in the tumor. However, genomic copy number evidence suggests an amplification leading to four copies of the RNase P control gene, suggesting that these dPCR results are likely false positives. Excluding these likely false positive calls from patient K, all of the retained and lost alleles were identified correctly by both DASH and LOHHLA (100% sensitivity and 100% specificity). To the best of our knowledge, the patient-specific dPCR presented here represents the first allele-specific genomic HLA LOH validation assay.

### Quantitative immunopeptidomics is limited as a validation approach

Mechanistically, HLA LOH is expected to effectively reduce the neoantigen load by eliminating surface presentation of neoantigens that would bind to specific HLA alleles. This mechanistic link has been demonstrated with organoids^21^, but it has not been shown in complex patient tumor samples. Thus, we sought to measure the quantitative changes in peptide presentation between adjacent-normal samples predicted by DASH to have or not have HLA LOH (**Figure 4A**). We performed quantitative immunopeptidomics with TMT labeling on three samples without any HLA LOH and four samples with predicted HLA LOH (**Supp. Table 6**). Across the seven samples, we found robust peptide yields (median ~5000 unique peptides), the expected length distribution (64.5% of peptides having nine amino acids), and a high percentage of observed peptides predicted to bind to at least one of the patient-specific alleles (**Supp. Figure 13A-B**). Then, we mapped the peptides to their predicted respective alleles using an MHC-Binding algorithm and looked for differences in peptide intensity for the peptides bound to lost, retained and homozygous alleles (**Supp. Figure 14-16**) ^27^. With all seven samples taken together, we found that peptides predicted to bind to lost alleles had reduced peptide intensity in tumor samples compared to normal samples for HLA-A and -B alleles (**Figure 4B**). HLA-C alleles, which have the lowest expression levels, did not show the same trend. However, the differences are very small and inconsistent across samples (**Supp. Figure 13C**). In fact, one of the samples that did not show the expected trend was previously validated as having HLA LOH with digital PCR (patient C), suggesting that either other factors also influence peptide presentation or the method is not robust. While a lack of consistently lower peptide intensities for lost alleles in tumor samples could be due to low tumor purities (**Supp. Figure 13D**) or variable differences in allele-specific expression (**Supp. Figure 13E**), we wanted to ensure that the method was robust to variabilities in immunoprecipitation and the limit of detection using mass spectrometry. Thus, we performed the same quantitative immunopeptidomics experiment with the CRL-5911 paired cell line used in the previous experiments. Unfortunately, the tumor cell line had very low expression of MHC and ten-fold higher peptide intensity values in the normal cell line than the tumor cell line. Due to these limitations, further work will be needed to functionally demonstrate the impact of HLA LOH on peptide presentation in patient tumor samples.

**Figure 4.**
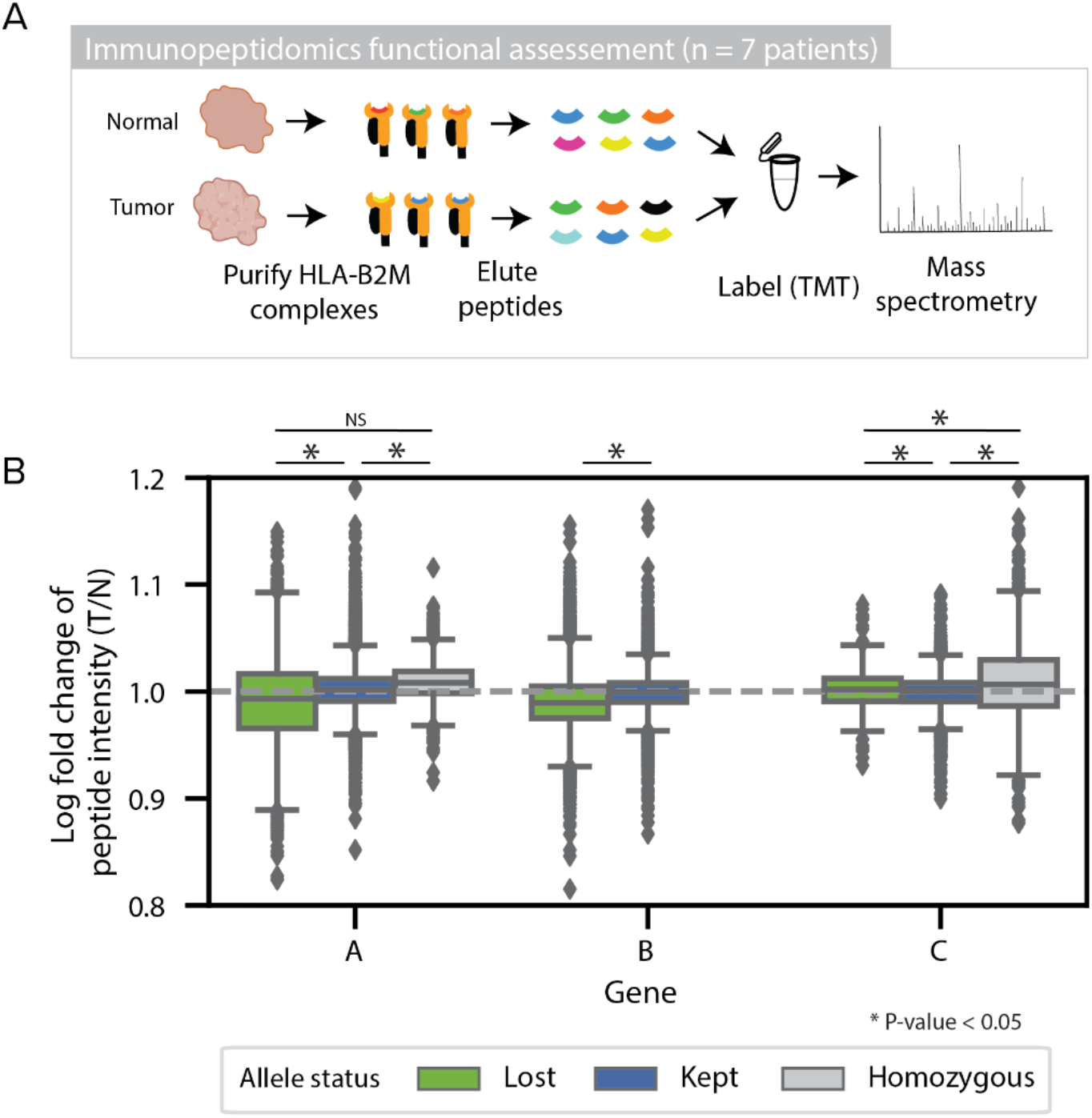
Functional immunopeptidomic validation. (**A**) Functional immunopeptidomic validation was performed using paired tumor and adjacent normal fresh frozen samples. HLA-B2M complexes were purified from each sample and peptides were gently eluted. Peptides from each sample were labeled with TMT tags and measured using quantitative mass spectrometry. (**B**) Box plots showing the log2 fold change of peptide intensity between lost, kept, and homozygous alleles across all patients. Statistical significance was assessed using a two-sided student T-test.

### HLA LOH is pervasive across tumor types and exhibits signs of immune pressure on tumors

To assess the pervasiveness of HLA LOH as a potential immune escape mechanism, we applied DASH to 611 tumors across 15 tumor types (see Methods). The tumors from the patients were predominately stage II and III, with a median tumor purity of 31% and a median MATH score (tumor heterogeneity) of 32.7 (**Supp. Figure 17A-C**). The fraction of patients with at least one incidence of HLA LOH ranged from 40% of patients in head and neck squamous cell carcinoma (HNSCC) to 4% of patients in liver cancer (**Figure 5A, Supp. Table 7**). We found a similar incidence of HLA LOH in non-small cell lung cancer adenocarcinoma (NSCLC-A) as compared to a previous study (24% and 29%, respectively), but found a significantly lower incidence in non-small cell lung cancer squamous cell carcinoma (NSCLC-SCC) (28% and 61%, respectively) ^22^. Of note, the overall frequency we report is similar to another study with a very large cohort size^14^. Moreover, we show that patients with HLA LOH often lose all three genes and more rarely lose only one gene or two genes (76.4%, 21.1% and 5.5% of patients, respectively; **Figure 5B**) and that HLA-A and -B are preferentially lost compared to HLA-C (**Supp. Figure 17D**). While some specific alleles appear to be lost much more frequently than other alleles, this trend is likely due to low starting frequencies of these alleles (**Supp. Figure 17E**).

**Figure 5.**
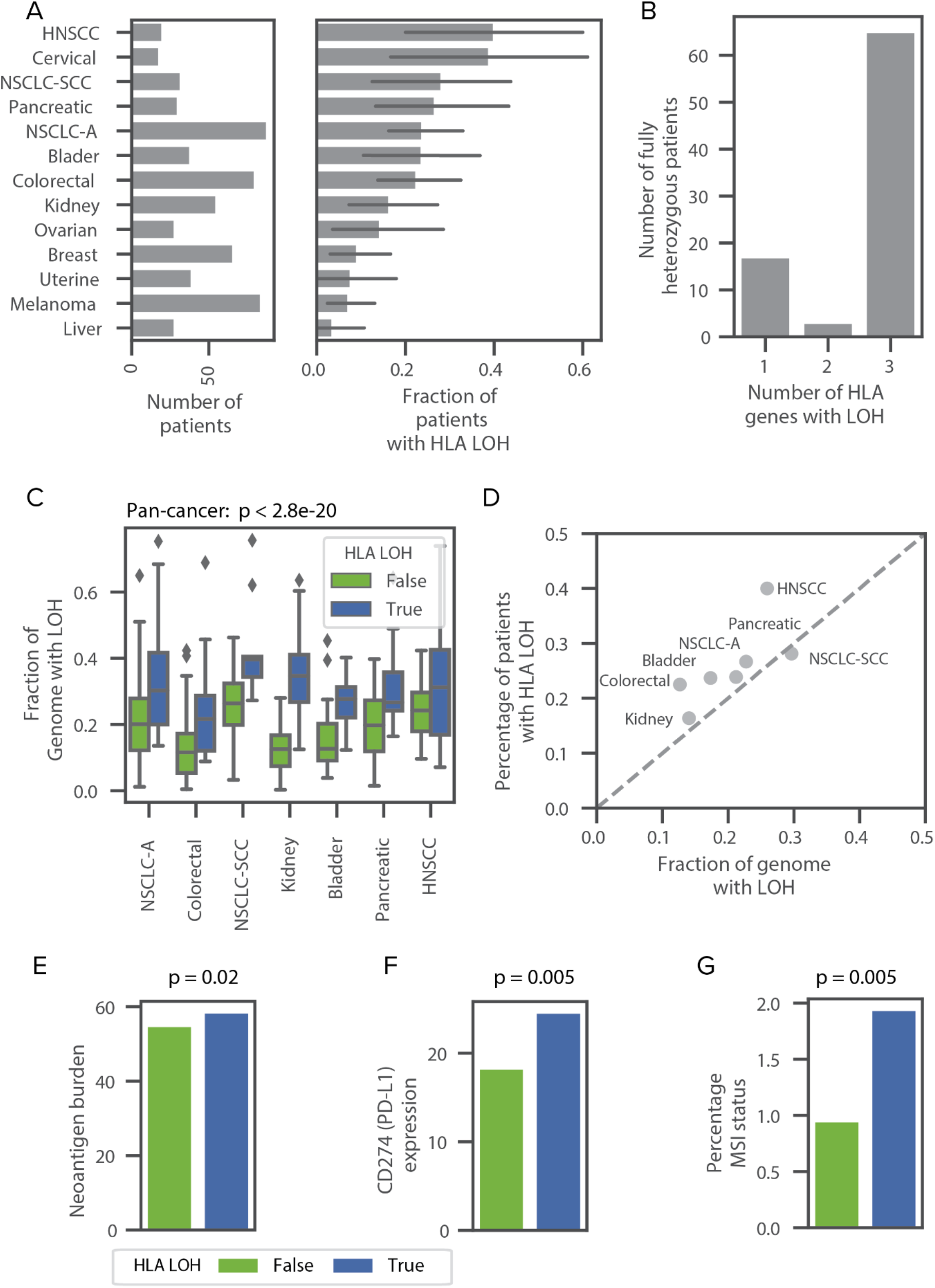
Widespread impact of HLA LOH across tumor types. (**A**) Bar plots denoting the number of patients and the frequency of HLA LOH in each tumor type cohort. Only cohorts with at least 10 patients are shown. (**B**) A bar plot showing the number of patients with 1, 2 or 3 genes impacted by HLA LOH. Only patients that are fully heterozygous across HLA-A, -B and -C are shown. (**C**) Boxplots showing the distribution of the fraction of each genome impacted by LOH. Each tumor type is divided into patients with HLA LOH and without HLA LOH. Only tumor types with at least 10 patients impacted by HLA LOH are shown. Statistical analyses are performed with two-sided Mann Whitney U tests and are Bonferroni corrected. (**D**) A scatter plot showing the relationship between the average fraction of the genome impacted by LOH and the frequency of HLA LOH in each tumor type. The grey dashed line denotes x=y. (**E-G**) Bar plots denoting the average difference for patients without HLA LOH (green) and patients with HLA LOH (blue) for (**E**) neoantigen burden, (**F**) CD274 (PD-L1) expression, (**G**) percentage of microsatellite sites with instability.

Though high frequencies of HLA LOH in specific tumor types are of interest due to impairment of the antigen presentation pathway, the high frequencies alone do not necessitate an evolutionary advantage of the LOH event. For example, we found that patients with HLA LOH have significantly higher estimated rates of LOH across their genome, suggesting that some LOH in the HLA region may happen by chance (pan-cancer p=2.8e-20 with the null hypothesis that patients with and without HLA LOH would have the same rate of LOH across their genome, **Figure 5C**). To investigate if HLA LOH frequencies across cancer types would occur by chance, we compared the average estimated rate of LOH across the genome with the frequency of HLA LOH in a given tumor type cohort. If LOH was randomly occurring in the HLA region, we would expect to observe the rate and frequency to be similar. However, we see that all tumor types individually and collectively have a higher frequency of HLA LOH than genome-wide LOH (**Figure 5D, Supp. Figure 18A**). While this difference is small for some tumor types, we observed that colorectal cancer and HNSCC had substantial enrichment of HLA LOH, suggesting that HLA LOH may provide a greater evolutionary advantage in these tumor types than others. Alternatively, HLA may be more prone to deletion than the rest of the genome.

Since HLA LOH impacts the ability of a tumor cell to present antigen on the cell surface for recognition by the immune system, we reasoned that tumors with a greater ability to display neoantigens would be under higher selective pressure to incur HLA loss. Considering patients with high HLA evolutionary diversity (HED) are able to present a larger immunopeptidome and respond better to checkpoint inhibitors^12^, we explored if they were more susceptible to HLA loss. However, we did not see this trend in our data (**Supp. Figure 18B**). Furthermore, high mutation rates and neoantigen burdens may also present pressure for cells to lose HLA. Indeed, we found correlations between mutation burden and HLA LOH pan-cancer (p=0.006 with the null hypothesis that patients with and without HLA LOH would have the same distribution of mutation burdens, **Supp. Figure 18C**) and between neoantigen burden and HLA LOH pan-cancer (p=0.02 with the null hypothesis patients with and without HLA LOH would have the same distribution of neoantigen burdens; **Figure 5E, Supp. Figure 18D**). Interestingly, we did confirm the “goldilocks effect” for mutation burdens with intermediate mutation burden tumors having the highest rate of HLA LOH ^19^, suggesting very high mutation rates may pose negative selection against HLA LOH (**Supp. Figure 18E**). Furthermore, we also corroborate correlations between HLA LOH and CD274 expression (PD-L1) and microsatellite instability (MSI) status (p=0.005 and 0.005, respectively with the null hypotheses that patients with and without HLA LOH would have the same distribution of CD274 expression and MSI status, **Figure 5F-G, Supp. Figure 18F-G**). The relationship between MSI status and HLA LOH suggests that some proportion of the HLA LOH events may be driven by genomic instability instead of immune pressure. We further hypothesized that HLA LOH may exert selective pressure on the mutational process as tumors develop. Indeed, we do observe that more neoantigens are predicted to bind to lost HLA alleles than their homologous counterparts (Wilcoxon rank sum test, p=0.01 with the null hypothesis that the same distribution of neoantigens would be predicted to lost and retrained HLA alleles), suggesting that HLA LOH is a key mechanism for the selective exposure of antigen to the immune system.

## Discussion

Accurate detection of HLA LOH is critical for its use as a biomarker for cancer immunotherapy. Although allele-specific alignment improves current heuristic methods ^22^, variability in exome capture across alleles and the relatively short sequences of the HLA genes introduce additional hurdles for identifying changes in copy number. Moreover, the field lacks methods to accurately and comprehensively determine the limit of detection, sensitivity and specificity of HLA LOH detection algorithms.

To address these limitations, we have implemented the first machine learning approach for HLA LOH detection, incorporating novel features that result in improved performance. The novel features include (1) a modified b-allele frequency equation to account for differences in probe capture, (2) consistency of sequencing depth to capture variance across the allele, (3) total sequencing depth ratio to detect hyper amplifications and (4) information from the regions surrounding the HLA genes because the vast majority of HLA LOH events are large deletions. Though DASH was trained using sequencing data from the ImmunoID NeXT Platform, our downsampling experiment demonstrates that DASH can be applied to datasets with lower sequencing depths with a limited decrease in performance.

To address the lack of well developed validation approaches for HLA LOH, we developed two methods to comprehensively evaluate algorithmic performance. Using *in silico* cell line mixtures, we utilized DASH’s high sensitivity and specificity across a range of tumor purities and sub-clonality levels and demonstrated superior ability to detect sub-clonal HLA LOH events than other available tools. To assess performance in actual tumor samples, we developed a novel PCR-based, allele-specific approach with sensitivity to detect sub-clonal HLA LOH. Of note, the scalability, sensitivity and allele-specific nature of this approach lends credibility to its potential clinical application. Movever, we tested the applicability of quantitative immunopeptidomics to observe the functional impact of HLA LOH. While we saw some significant differences in peptide intensities, more work must be done to test the sensitivity and robustness of this method as well as measure the impact of other biological factors, including infiltrating immune cells, variable expression or translation and peptide loading conditions, that may disrupt the signal.

Our HLA LOH prevalence data shows that a large percentage of patients are impacted by HLA LOH in several tumor types. Though NSCLC was previously reported to have a high incidence of HLA LOH ^22^, we also identified a large fraction of HLA LOH in cervical cancer (38%) and HNSCC (40%). NSCLC and HNSCC are also known for having a high mutational burden; however, we observe only 14% of patients with HLA LOH in melanoma, which also has a high mutational burden ^28^. Cervical cancer is strongly associated with human papillomavirus (HPV), which may play a role in the high frequency of HLA LOH. Of note, we observed a lower frequency of HLA LOH in NSCLC-SCC than one previously published study ^22^ and a similar frequency to another study ^14^. Differences in tumor stage, clonality and treatment status may contribute to these differences. Larger cohorts and comparisons of tools within the same dataset are needed to understand all underlying covariates. Our analyses also revealed that patients tend to lose more than one HLA-allele at a time, potentially having stronger implications on tumor evolution.

Our study has several limitations. Larger cohorts of patients will allow for more scalable validation and increase confidence in the accuracy of the manual annotations used for training. Further, we have provided a proof-of-concept for the detection of the functional impact of HLA LOH with quantitative immunopeptidomics. However, more work must be done to evaluate the robustness of this approach. Finally, detecting HLA LOH alone does not capture all allelic-imbalance driven immune escape mechanisms. Methods to detect significant amplification of a single allele, potentially allowing the drowning out peptides from the homologous allele, and allele-specific expression are also needed.

In conclusion, DASH is a novel, validated HLA LOH algorithm. We demonstrated DASH’s ability to sensitively detect sub-clonal events in samples with low tumor purity, enabling comprehensive profiling of widespread HLA LOH across tumor types.

## Methods

### Patient data used for model training and testing

To train our HLA allele-specific deletion algorithm, we collected tumor and adjacent normal or blood normal samples from 279 patients across 15 different tumor types (a subset of the 611 patients used in later sections). These samples were purchased from various biobanks and were collected under IRB-approved protocols. For each patient, paired FFPE or fresh frozen samples were profiled using Personalis’ ImmunoID NeXT Platform; an augmented exome/transcriptome platform and analysis pipeline, which produces comprehensive tumor mutation information, gene expression quantification, neoantigen characterization, HLA (typing and mutations) and tumor microenvironment profiling. Each component is outlined in a separate section.

### ImmunoID NeXT - Whole exome sequencing

For all samples in this study, whole exome library preparation and sequencing were executed as previously described ^29^. DNA from tumor and PBMC/adjacent sample were used to construct whole-exome capture libraries using two whole-exome sequencing (WES) capture kits: Agilent SureSelect Human All Exon v5 plus untranslated regions and Agilent SureSelect Clinical Research Exome, as recommended by manufacturers. Modifications were made to the protocols to yield an approximately 250 bp average library insert size. Moreover, a KAPA HiFi DNA Polymerase (Kapa Biosystems) was used instead of Herculase II DNA polymerase (Agilent). Finally, sequencing was performed using either HiSeq 2500 or NovaSeq instrumentation (Illumina) at 20 Gigabases (G) sequencing depth for normal samples and 35G sequencing depth for tumor samples (G = (2 * # of reads * read length) / 1000000000; used instead of sequencing depth due to variable sequencing depths). We observed over >300x sequencing depth exome-wide for >20,000 genes and >1000x sequencing depth for the exons of boosted regions consisting of over 500 cancer-associated genes, including HLA-A, -B and -C.

### ImmunoID NeXT - Alignment

All DNA and RNA reads are aligned to the hs37d5 reference genome build (BWA v0.7.12), in accordance with the Personalis ACE Cancer Exome Analysis Pipelines. The pipelines follow the best practice guidelines as recommended by the Broad Institute. The Picard toolkit is used for duplicate read removal (Picard v1.74). The Genome Analysis toolkit (GATK) is used to correct base quality scores and improve sequence alignment.

### ImmunoID NeXT - HLA typing, HLA typing validation and HLA somatic mutations

To perform next generation sequencing-based HLA-typing, the blood or adjacent normal sample from a patient was used. HLA typing was performed using the ImmunoID NeXT Platform. HLA types were calculated up to 6 digits. To validate the HLA typing approach, we performed blinded typing on 15 samples with known HLA genotypes. Ten of the samples were procured from ASHI (American Society of Histocompatibility and Immunogenetics) and consisted of 2.5mL of whole blood preserved in acid citrate dextrose. Five of the samples were procured from CAP (College of American pathologists) and consisted of 2.0mL of whole blood preserved in citrate phosphate dextrose or citrate phosphate dextrose-adenine. Genomic DNA was extracted from the blood and sequenced using the ImmunoID NeXT Platform. The results are shown in **Supp. Table 1**. For the HLA somatic mutation calling, the tumor and normal data were integrated with Polysolver ^8^ using default parameters (v1.0.5, arguments: reference - hs37d5, data type - STDFQ).

### ImmunoID NeXT - Tumor copy number alterations, purity and ploidy estimates

Sequenza ^15^ was run with default parameters on each sample (v1.0.3). Allele-specific copy number alterations were obtained from *_segments.txt output file. Tumor purity (or cellularity) and tumor ploidy were extracted from the first line of the *_confits_CP.txt output file.

### Creation of HLA allele database

The HLA allele database was created with an imputation approach similar to that described in ^8^. The database was initialized with the multiple sequence alignment (MSA) format from IMGTv312 ^23^, the same version used by HLAssign for HLA typing. Then, we used the cDNA file to impute exons in alleles and incompletely sequenced alleles with a reference allele that had protein-level identity as defined by identical 4 digit nomenclature. If no such allele existed, we used a reference from the same HLA subtype, as defined by identical 2 digit nomenclature. If there were multiple options with identical 2 digit nomenclature, we then used the first allele listed in the MSA. To impute the intronic regions of each allele, we used the same approach with the gDNA file. The full length genomic sequences of each allele were imputed by assembling exons from the cDNA imputation set and the introns from the gDNA imputation step.

### Feature assembly for DASH

For each sample, the patient’s HLA alleles are used to create a patient-specific HLA reference. Our HLA typing aggregates all reads that could map to the HLA region using a 30 base pair seed. We used BWA to map all of these reads to the patient-specific HLA reference ^30^. Since ImmunoID NeXT has high sequencing depth of the HLA region (average of 1586 sequencing depth over exonic regions; average of 587 sequencing depth over each intronic and exonic position after post-processing), we were able to be very stringent about the reads we chose to map. We excluded any reads that had soft clipping for more than 20% of their total length. Furthermore, we required exact sequence matches, discarding any reads that contained mismatches in order to attain the highest quality sequencing depth information. However, if we detected a somatic mutation within the HLA alleles, we lifted the stringency to allow reads with a single mismatch. Sequencing depth at each position of each patient-specific HLA allele was calculated using Samtools ^31^.

Next, we aligned the patient-specific homologous alleles to determine positions of difference between the alleles. We detected both single nucleotide variants (SNVs) and indels in the alignment. We only considered these points of difference in our sequencing depth related features because all other positions are impacted by the presence or absence of the homologous allele. Importantly, only the first position of each indel was considered to ensure SNVs were appropriately weighted. Alleles with fewer than 5 positions of difference between them were considered to be homozygous.

From the sequencing depth data, we calculated the following allele-specific features:

1. *Adjusted b-allele frequency (BAF):* At each position of mismatch, the b-allele frequency is calculated for the tumor and normal sample separately. Then, the tumor BAF is divided by the normal BAF. Using the normal sample to adjust the BAF is critical because there is variability in the probe capture of each specific allele. To consolidate the ratios into a single feature, the allele references are broken into bins of 150 base pairs in length. The absolute value of the median adjusted-BAF is calculated for each bin. Then, the median value across all bins is used as the feature. The adjusted b-allele frequency feature has a lower bound of 0, with larger numbers denoting a higher chance of LOH in the HLA gene (**Supp. Figure 1A**). For the following equation, BAF represents b-allele frequency, T represents tumor and N represents normal:

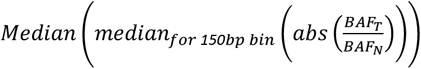
2. *Allele-specific (AS) sequencing depth ratio*: At each position of mismatch between the homologous alleles, the ratio of sequencing depth in the tumor sample to the sequencing depth in the normal sample is calculated for each allele. Each value is normalized by the exome-wide number of tumor reads divided by the exome-wide number of normal reads. Thus, despite variability in sequencing depth between each run, the expected allele-specific sequencing depth ratio is one if there is no copy number variation. Then, for each bin, the median sequencing depth ratio is calculated for each allele and the lower value amongst the two alleles is considered for that bin. Finally, the median value across all of the bins used as the feature. The allele-specific sequencing depth ratio has a lower bound of 0, with an expected value of 1 if there is no copy number variation. Lower allele-specific sequencing depth ratios suggest a high probability of LOH in an HLA gene (**Supp. Figure 1B**). For the following equation, AD represents A-allele sequencing depth, BD represents B-allele sequencing depth, R represents total reads across the exome, T represents tumor and N represents normal:

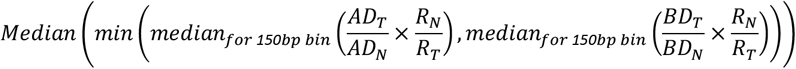
3. *Consistency of sequencing depth:* An allele that has consistently lower sequencing depth across all mismatch positions is more likely to be a real case of HLA LOH. Thus, we give each allele a 0 or 1 for each bin depending if it has lower or higher sequencing depth compared to its homologous allele. If the allelic sequencing depth of a bin cannot be determined (no mismatch sites), each allele is given a value of 0.5. Then, we take the average across all bins for each allele and assign the higher average to be the value for the feature. The consistency of the sequencing depth feature ranges from 0.5 to 1 with values closer to 1 representing a higher likelihood of HLA LOH (**Supp. Figure 1C**). For the following equation, AD represents A-allele sequencing depth, BD represents B-allele sequencing depth, T represents tumor and N represents normal:

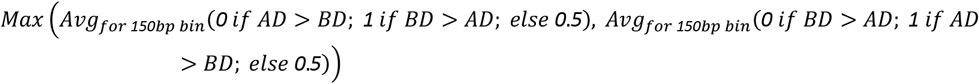
4. *Total sequencing depth ratio:* If a heterozygous pair of alleles has a high combined sequencing depth, allelic imbalance may be driven by a large amplification in an allele rather than a deletion in an allele. At each position of mismatch between the homologous alleles, the ratio of sequencing depth in the tumor sample to the sequencing depth in the normal sample is calculated for each allele. Then, the sum of both alleles is taken as the value representing each 150 base pair bin. Finally, the median across the bins is used as the total sequencing depth ratio feature. The total sequencing depth ratio has a minimum of zero, with higher values tending toward genes without HLA LOH (**Supp. Figure 1D**). For the following equation, AD represents A-allele sequencing depth, BD represents B-allele sequencing depth, R represents total reads across the exome, T represents tumor and N represents normal:

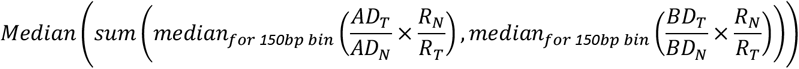

In addition, we included patient-specific features:

1. *Tumor purity:* The tumor purity feature is obtained from the Sequenza tumor purity estimation. The value ranges from 0.1 to 1, with 0.1 being the least pure tumor and 1 being the most pure tumor.
2. *Tumor ploidy:* The tumor ploidy feature is obtained from the Sequenza tumor ploidy estimation. The values are whole integers that are greater than or equal to one.

Finally, we also included a whole-exome feature:

1. *Deletion of flanking regions:* Since most instances of HLA LOH are due to large deletions, we created a feature representing deletions in the flanking regions of each gene to capitalize on information from a larger number of variable sites. We use the b-allele deletion calls from Sequenza and make a flanking region deletion call if there is a deletion within 10,000 base pairs in either direction of the gene. This feature is binary, with 0 representing a deletion.

### Manual labeling of HLA LOH in training dataset

The heterozygous HLA genes for a set of 279 patients were visualized and manually annotated for HLA LOH. Of the 279 patients, 12 patients were excluded based on poor data quality or the ambiguous natures of their HLA LOH status. As a result, 267 patients with 720 heterozygous genes were manually labeled for training. Heterozygous genes were visualized as shown in **Supp. Figure 2** and labeled based on the following criteria: 1) the tumor sequencing depth plots were compared to the normal sequencing depth plots to look for a consistent decrease in one allele in the tumor data; 2) the tumor b-allele frequencies were compared to the normal b-allele frequencies to look for a consistent decrease or increase across the vast majority of the mismatch sites; 3) the exome-wide normalized, allele-specific sequencing depth ratios were assessed for a significant depletion of a single allele below one; 4) the a- and b-allele copy numbers in the regions flanking the gene were assessed, with b-alleles of 0 suggesting HLA LOH; 5) criteria 1-3 were considered in light of the tumor purity, with lower tumor purities requiring less substantial differences; 6) criteria 1-3 were considered in light of the tumor ploidy, with high tumor ploidies lending higher weight to the flanking regions due to the instability of the allele-specific sequencing depth ratios. The manual labeling was updated once during development after receiving the first round of digital PCR results to further weight the flanking regions in cases with high amplification of a single allele in order to increase specificity on these samples with high allelic imbalance.

### Training the DASH algorithm

A set of 720 heterozygous genes were collected from 279 patients across multiple tumor types. Since scalable gold standard HLA LOH labeling is cost and labor prohibitive, we visualized all features described above for each heterozygous gene and manually labeled each case of HLA LOH. To evaluate the performance of DASH, we used a total of 720 heterozygous genes as input for 10-fold cross-fold validation. For the final method, we used all 720 heterozygous genes in training. Then, for both the evaluation and final method, we employed an XGBoost algorithm to learn how to detect HLA LOH in each pair of alleles from the features described above. The XGBoost algorithm was trained with the following parameters: loss function - binary logistic; max depth - 5; eta - 0.3; subsample - 0.5; min child weight - 2; max delta step - 0; number of estimators - 100. We compared the XGBoost algorithm to three other machine learning algorithms: linear regression, support vector machine and k nearest neighbors. To improve the specificity of DASH on samples with low tumor purity, we also performed a secondary check to ensure that the allele-specific ratio of the lost allele is < 0.98 and that the adjusted-BAF is > 0.02. If HLA LOH was detected by the algorithm, the allele with the lower sequencing depth was labeled as deleted. Though rare, we observed a small minority of cases with a bi-allelic deletion. If the algorithm detects HLA LOH and the allele with higher sequencing depth has an allele-specific sequencing depth ratio below 0.5 for at least 25% of the bins, both alleles are labeled as deleted. In this manuscript, DASH was run with default parameters as outlined in the supplementary code and GitHub (https://github.com/Personalis-DASH/DASH.git).

### Sequencing depth downsampling analysis

In order to assess DASH’s performance with lower exome sequencing depth data, a computational downsampling method was used. We derived the HLA sequencing depth of standard exomes from whole exome data from dbGaP (NCBI database of Genotypes and Phenotypes, RRID:SCR_002709) (study accession: phs000452.v3.p1) ^24,25^. We then performed downsampling on each sample to sequencing depths at the HLA region of 10X, 30X, 50X, 100X, and 200X to represent the range of potential whole exome sequencing depths. We calculated the number of reads needed for each sequencing depth as the sum of the sequencing depth values for homologous alleles from all reads that fit DASH’s strict alignment requirements. Once the number of reads needed to ascertain the desired sequencing depth was calculated, we used ten different number seeds to generate ten uniquely downsampled BAM files for each sample and then used these BAM files as input to DASH. All non-allele-specific features were held constant with the original sample. Using the labeled 720 heterozygous alleles, we analyzed F1-scores across all seeds for each desired depth as well as F1-scores across all seeds for each desired depth, split up by a tumor purity inclusion threshold, where each increasing subsequent threshold value (0.2, 0.4, etc.) includes the samples with tumor purity up to the stated tumor purity level.

### Cell line based limit of detection analysis

To assess the limit of detection of DASH across varying tumor purities and clonalities, we identified a tumor-normal paired lymphoblast cell line (NCI-H2009). In NCI-H2009, HLA-A is homozygous while both HLA-B*51:01 and HLA-C*15:02 alleles are deleted. We deeply sequenced the tumor and normal cell lines to 50G and 30G, respectively. In order to stimulate a realistic sequencing depth, we downsampled the normal data to reflect 20G. To create tumor data of decreasing purity, we mixed increasing proportions of normal reads with decreasing proportions of tumor reads (**Supp. Table 1**). The combined normal and tumor reads always sum to an average of 35G sequencing depth to represent the tumor sample. All combinations of normal and tumor sub samples were taken from the same sequencing runs and performed in replicates of 10 using the seqkit library. To simulate lower sub clonality, the whole exome features and the allele-specific features were determined separately. The whole exome features were derived from a whole exome read mixture that reflected stated tumor purity with the representative percentage of tumor reads with the percentage of normal reads. For the allele-specific features, the proportion of tumor reads used in the mixture was the product of stated tumor purity and clonality level. Samples without HLA LOH were simulated by only including normal reads in the tumor sample and artificially increasing the estimated tumor purity to reflect the desired range. These runs were used to estimate specificity.

### Comparison of performance to LOHHLA

To compare DASH against a known and tested tool for detection of LOH in HLA alleles, we executed LOHHLA (Loss of Heterozygosity in Human Leukocyte Antigen), developed by McGranahan et al., to call HLA LOH in our test dataset and our *in silico* cell line dilutions ^22^. As input, LOHHLA requires tumor/normal matched BAM files. Thus, we aligned our normal and tumor FASTQ files to hg19 with BWA, specifying the retention of unmapped reads. To determine HLA LOH calls, we used the cutoffs specified in the LOHHLA manuscript: copy number of an alternate allele is < 0.5 and the p-value related to this allelic imbalance is < 0.01. Two samples that were correctly predicted by DASH produced failure errors with the LOHHLA and were held out of the LOHHLA performance analysis (DNA_ILS43894PT2 and DNA_32652).

### Allele-specific digital PCR validation

To experimentally validate allele-specific HLA LOH in samples, we designed patient-specific primers and probes and tested for depletion of allele-specific DNA with digital PCR (dPCR). Since each patient has a unique set of up to 6 HLA class I alleles, patient-specific primers and probes must be designed for each patient. These primers and probes must bind with high specificity to each allele of interest and discriminate against all other alleles and the rest of the genome. Due to the similarity of some homologous alleles, good primers and probes may not exist for all patients. We manually designed primers and probes for eleven homologous allele pairs with HLA LOH predicted by DASH from ten different patients and one cell line to maximize descrimination between alleles (**Supp. Table 4**). Furthermore, a probe targeting RNase P (RPP25) was also used to serve as an internal positive control. The HLA allele and RPP25 probes were assigned different fluorescence to allow multiplexing (FAM and VIC, respectively). H20 was used as a negative control sample.

To assess the efficiency of the primers and probes, dPCR was performed in triplicate on the DNA from the normal and tumor samples (excluding patient C, which was performed in duplicate). Four samples were from the training dataset (A, B, I, K) and the remaining seventeen samples were fully independent. To analyze the data, both the lost and retained allele will be normalized by the control gene to account for sample input variation. The primers and probes were deemed successful if the ratio of the HLA allele copies to the multiplexed RNase P copies was 0.5 in the normal sample because the HLA allele was expected to be haploid and RNase P is expected to be diploid. Then, for the primer designs that fit this requirement, the allele to RNase P ratio in the tumor DNA is compared to the allele to RNase P ratio in the normal DNA with a one-sided T-test (with the null hypothesis that the tumor would be greater than or equal to the normal) to determine if there has been a significant drop in the tumor. The variability of the dPCR measurements are assumed to be normally distributed. This test is performed for both the predicted retained allele and the predicted lost allele. Allelic imbalance is determined by measuring a significant difference between the predicted lost and predicted retained alleles in the normal DNA and the tumor DNA. Of note, this validation focuses on specific sections of each gene. Thus, it is not formulated to catch small focal deletions in a small portion of the gene.

### Quantitative immunopeptidomics validation

To assess the functional impact of HLA LOH on peptide presentation by MHC, we performed quantitative immunopeptidomics on one pancreatic, two distinct colorectal and four distinct lung tumor-normal paired fresh frozen samples (51 mg - 819 mg range) as well as one paired tumor-normal cell line (CRL-5911). The samples were homogenized, normalized for protein content between the tumor and normal and the clarified homogenates were applied to a pan-MHC-I antibody (W6/32)-linked immunoaffinity resin. The success of immunoprecipitation from the lysates was assessed using ELISA, by comparing the MHC concentration pre- and post-IP. MHC-associated peptides were gently eluted and collected. Eluted peptides from tumor and normal samples were labeled using 130C and 131N channels of a 10-plex tandem mass tag kit (ThermoFisher Scientific; lot# UK282322) according to the standard protocol and analyzed by LC-MS/MS in a single run for each pair, in high resolution HCD mode (Fusion Lumos). The immunopeptidomics datasets generated during and/or analyzed during the current study are available in the PRIDE repository with project accession PXD022323 ^32^.

The resulting raw files of all six samples were processed together by PEAKS. Peptide identification was performed using the standard PEAKS pipeline, i.e. *de novo* identification followed by database search. Parameters for the database search were as follows -- precursor mass tolerance: 10 ppm, fragment mass tolerance: 0.03 Da, protein database: uniprot sequences downloaded in April 2019, enzyme digestion: none, fixed modifications: carbamidomethylation of cysteine (+57.02 Da) and TMT10plex at all N-terminal amino acids and lysines (+2291.6), variable modifications: protein N-terminal acetylation (+42.0106) and oxidation of Methionine (+15.9949). Peptides were filtered at 1% FDR and reporter ions were quantified using the PEAKS Quant module. The list of quantified peptides were further filtered using an in-house script to increase the quality of calls by removing peptides that do not have expected TMT n-terminal or lysine modifications, peptides with low intensity (less 10E4 precursor ion intensity) and suspicious peptides with poly amino acids. Then, the intensities were log2 transformed and the data was median normalized. Finally, a fold change was calculated from the log2 transformation, with values less than 0 representing a depletion of peptide in the tumor sample and values greater than 0 representing enrichment of peptide in the tumor sample.

To assess overall changes in presentation between the normal and tumor samples, we compared the absolute value of the logarithm of the fold changes amongst the samples. Next, we estimated the peptide change for specific alleles. For each patient, we used an MHC presentation prediction algorithm (described below) to assign each peptide to an MHC allele. If a peptide was predicted to bind to multiple alleles, it was considered ambiguous and excluded from the analysis. If the only peptides predicted to bind to an allele were predicted to bind to more than one allele, these were included but the allele was marked with an asterisk. The logarithm of the fold change was visualized to assess the enrichment or depletion of peptides from particular alleles in the tumor sample or across a set of alleles or patients. The log2 transformed intensity values were compared with a two-sided Wilcoxon Rank Sum Test to assess the statistical significance of any enrichment or depletion. All comparisons with tumor purity use the tumor purity as estimated by Sequenza.

### Predicting HLA-associated neoantigens

MHC class I binding prediction was performed with SHERPA ^27^. The percentile rank threshold used in this manuscript is 0.1%. All peptide-allele combinations with ranks below this threshold are considered to bind.

### Pan-cancer HLA LOH frequency across tissue types

A total of 611 patients from across 15 tumor types were considered for analysis. Each patient had a tumor sample and a normal sample that were sequenced and analyzed on the ImmunoID NeXT Platform, as described above. Patients without HLA calls due to low coverage were excluded from the frequencies (n=2). A subset of these samples were used for training the DASH algorithm. DASH was run on each sample to predict the genes (HLA-A, -B and -C) that were impacted by HLA LOH. The frequencies of HLA LOH cooccurrence between multiple genes within a single patient were calculated based on a reduced cohort that only contained fully heterozygous patients.

### Genome-wide LOH analysis

As part of ImmunoID NeXT, copy number variation was estimated with Sequenza. For each region with variable copy number, if the estimated copy number of the b-allele was zero, the region was considered LOH. The total number of base pairs impacted by LOH were totaled for each patient and divided by the total number of base pairs across the exome (3.2 billion) to obtain the fraction of the genome with LOH. Though the fraction may be an underestimate due to limited sequencing depth of genomic regions without genes, we expect the underestimation to be consistent across patients.

### Sequence divergence analysis

The germline HLA-I evolutionary divergence (HED) score was calculated from the HLA alleles of each individual patient as described in ^12^. The HED score is intended to capture allelic sequence diversity for a patient’s HLA alleles, with a low score denoting low diversity and a high score denoting high diversity.

### Mutational and neoantigen load analysis

Mutational burdens were generated using tumor-specific genomic events (SNVs, indels, and fusions) of at least 5% allelic fraction that were verified using transcriptomic data. All potential neoepitopes (8-,9-,10- and 11-mers) were created for each mutation and tested for presentation as described above. If any 8-,9-,10- or 11-mers containing the mutation were predicted to bind to any of the patient-specific alleles, they are considered putative neoepitopes.

### Gene and HLA allele expression analysis

Alignment of whole-transcriptome sequencing was performed using STAR ^33^. Transcripts per million (TPM) were calculated to normalize the expression values and used to test for correlations with HLA LOH. HLA allele specific TPMs were estimated via kallisto 0.45.1 and customized transcriptomic references. Individualized transcriptomic references were generated by starting with the Ensembl version 103 / GRCh37 cDNA reference and replacing reference HLA sequences with individual-specific typed HLA allelic sequences based on IPD-IMGT/HLA 3.43.0 CDS reference sequences. These individualized references were then indexed by kallisto separately, followed by TPM quantification using kallisto’s EM algorithm to adjust for allelic sequence similarities both in the HLA locus and genome-wide.

### Microsatellite Instability analysis

The percentage of microsatellite sites with instability was determined using MSIsensor, a computational tool that queries sequencing reads from specific microsatellite sites from matched tumor-normal sequencing data ^34^.

## Author Contributions

Original concept - R.M.P and S.M.B.

Project supervision - S.M.B. and R.C.

Algorithmic design and evaluation - R.M.P., S.D., S.M.B. and S.V.Z.

In silico cell line analysis - R.M.P. and S.D.

PCR experimental design and analysis - R.M.P., D.M., and S.M.B

Immunopeptidomics data processing - D.M. and R.M.P.

Immunopeptidomics analysis - R.M.P. and D.M.

Pan-Cancer cohort collection - C.A.

Pan-Cancer cohort analysis - R.M.P. and S.D.

HLA typing validation experiment - G.B.

Figures - R.M.P. and S.D.

Manuscript - R.M.P., S.M.B, D.M., S.D., R.C., J.W., J.S., D.C., E.L.

## Acknowledgements

Digital PCR was conducted at the Genetic Resources Core Facility, Johns Hopkins Institute of Genomic Medicine, Baltimore, MD. The quantitative immunopeptidomics experiment was conducted at Cayman Chemicals. The primers were designed with DNA Software.

