## Supplemental Figures and Legends for "Validated machine learning algorithm with sub-clonal sensitivity reveals widespread pan-cancer human leukocyte antigen loss of heterozygosity"

### This PDF file includes:

Supplementary Figures 1-18

### Other Supplementary Materials for this manuscript include:

**Supplementary Table 1. HLA typing validation**

List of samples and HLA types used for HLA validation and concordance results of the experiment.

**Supplementary Table 2. Cell lines profiled for HLA LOH**

List of cell lines profiled by ImmunoID NeXT for HLA LOH, along with details about the cell lines used for the downsampling experiment and the specific HLA alleles that were deleted.

**Supplementary Table 3. Cell line downsampling**

Detailed overview of purity, clonality, number of normal reads, number of tumor reads, percentage of tumor reads and replicates for the cell line downsampling analysis.

**Supplementary Table 4. Primer designs**

Primer and probe sequences paired with sample information, lost gene, LOH status prediction.

**Supplementary Table 5. Raw digital PCR results**

FAM and VIC copies for each replicate and allele of each (A) cell line dilution and (B) patient.

**Supplementary Table 6. Peptides and log fold change from quantitative immunopeptidomics**

Processed quantitative immunopeptidomics data for each patient. Columns include the peptide and the log fold change intensity.

**Supplementary Table 7. HLA LOH frequencies by tumor type**

Total number of patients and frequency of patients with HLA LOH. Data behind Figure 5A.

**Supplementary Code.** Scripts for reproducing the analysis and models.

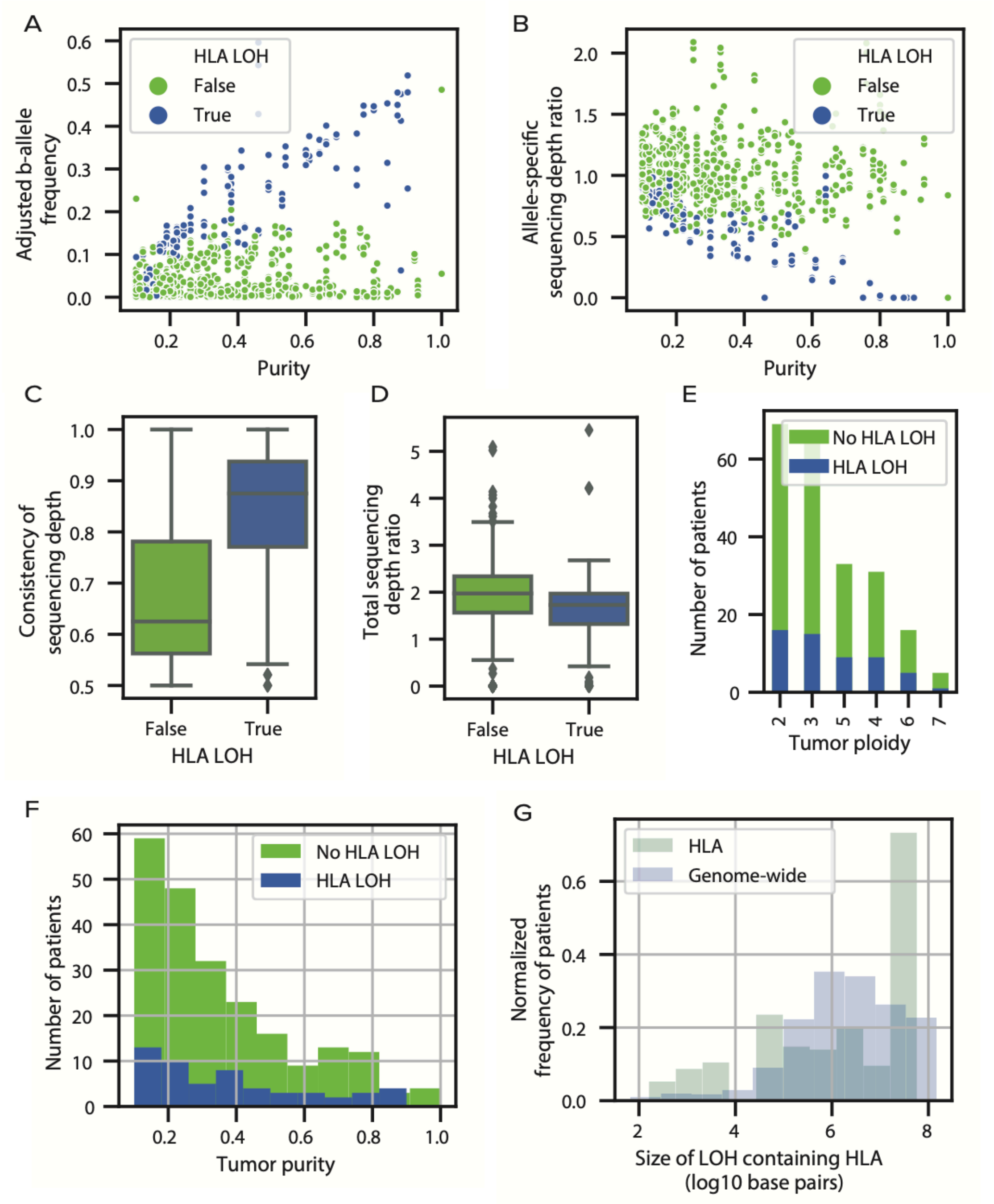


**Supplementary Figure 1. Overview of features in training data set.** (**A**) A scatter plot showing the relationship between Adjusted B-allele Frequency and tumor purity. Genes with HLA LOH are shown in blue and genes without HLA LOH are shown in green. (**B**) A scatter plot showing the relationship between allele-specific sequencing depth ratio and tumor purity. Genes with HLA LOH are shown in blue and genes without HLA LOH are shown in green. (**C-D**) Boxplots showing the difference in distribution between (**C**) consistency of sequencing depth and (**D**) total sequencing depth for patients with and without HLA LOH. (**E-F**) Histograms showing the distributions of tumor (**E**) ploidy and (**F**) purity for patients with and without LOH. Genes with HLA LOH are shown in blue and genes without HLA LOH are shown in green. (**G**) A histogram showing the distribution of LOH sizes that contain HLA genes across all patients.

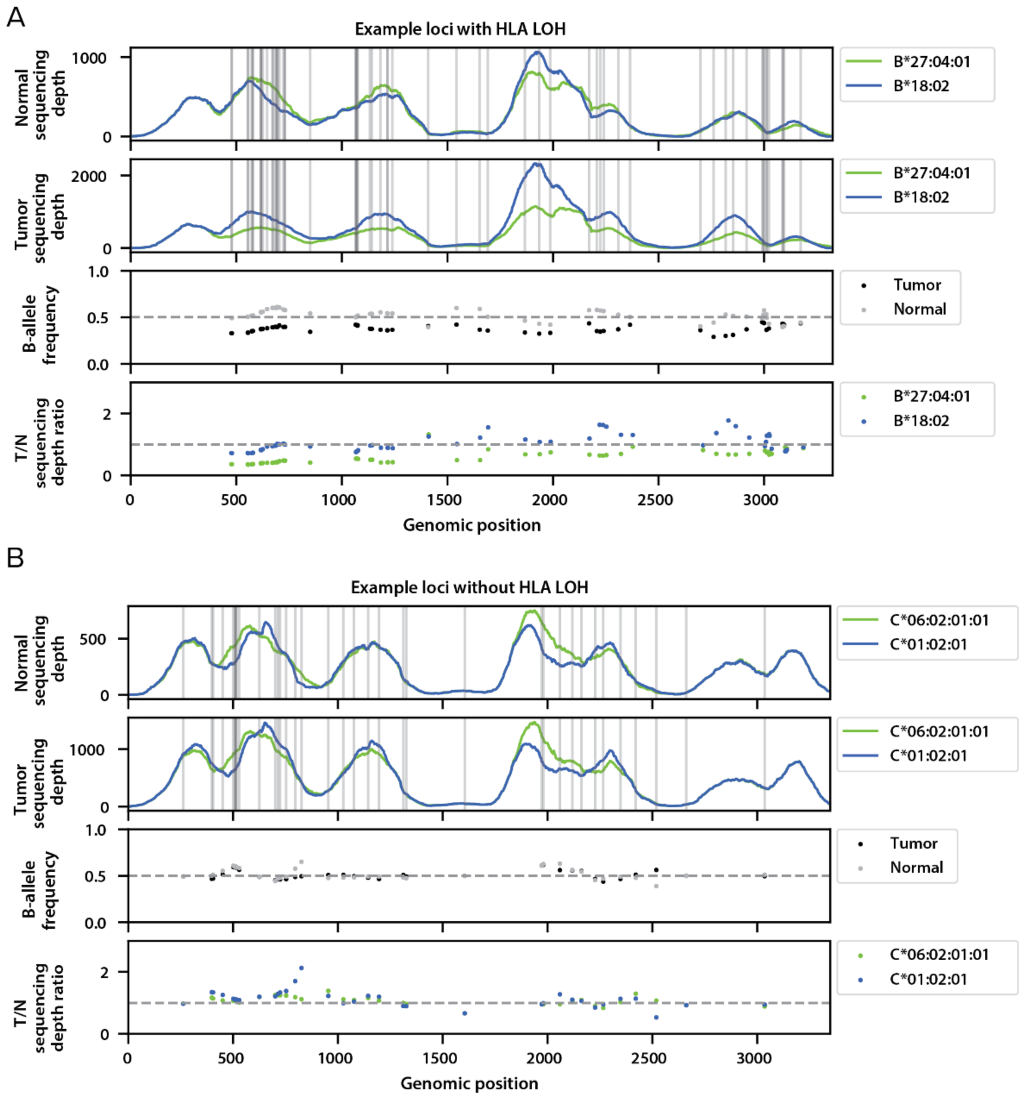


**Supplementary Figure 2. Overview of features in manual annotations.** (**A**) Example of the feature output of a loci containing a manually annotated HLA LOH event. Blue and green denote the different alleles for a specific loci for normal sequencing depth, tumor sequencing depth, and tumor / normal (T/N) sequencing depth ratio. Black and white denote the tumor or normal B-allele frequency respectively. Normal sequencing depth, Tumor sequencing depth, B-allele frequency, and T/N sequencing depth ratio were all used to determine if a sample was manually annotated with HLA LOH. **(B)** Example of features of a loci not containing a manually annotated HLA LOH event.

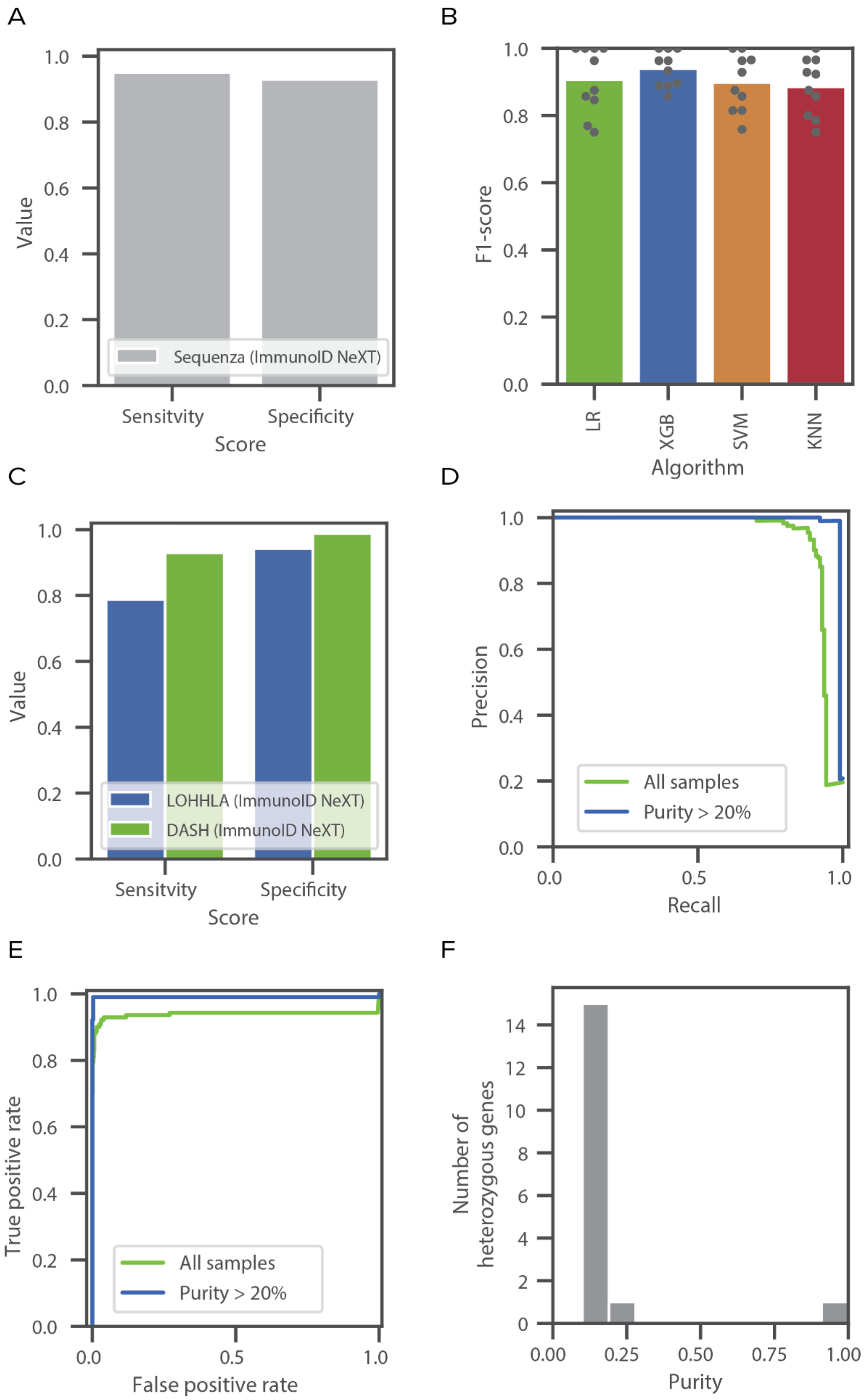

A
1.0
0.8
0.6
0.4
0.2
0.0
Value
Sequenza (ImmunoID NeXT)
Sensitvity
Specificity
Score
B
1.0
0.8
0.6
0.4
0.2
0.0
F1-score
LR
XGB
SVM
KNN
Algorithm
C
1.0
0.8
0.6
0.4
0.2
0.0
Value
LOHHLA (ImmunoID NeXT)
DASH (ImmunoID NeXT)
Sensitvity
Specificity
Score
D
1.0
0.8
0.6
0.4
0.2
0.0
Precision
All samples
Purity > 20%
0.0
0.5
1.0
Recall
E
1.0
0.8
0.6
0.4
0.2
0.0
True positive rate
All samples
Purity > 20%
0.0
0.5
1.0
False positive rate
F
14
12
10
8
6
4
2
0
Number of
heterozygous genes
0.00
0.25
0.50
0.75
1.00
Purity

**Supplementary Figure 3. Performance of DASH. (A)** Bar plot showing the sensitivity and specificity of using Sequenza to detect HLA LOH in ImmunoID NeXT samples. **(B)** Bar plot of F1-scores of different classifier machine learning algorithms using a 10-fold cross validation method. Dots represent the performance of the 10 individual algorithms. Abbreviations are as follows: LR - linear regression, XGB - XGBoost, SVM - support vector machine, KNN - K nearest neighbor. **(C)** Bar plots showing the sensitivity and specificities scores across ImmunoID NeXT samples between LOHHLA (blue) and DASH (green). **(D)** Precision-recall curve for DASH across all samples in green and all samples with a tumor purity > 20% in blue. **(E)** ROC-curve showing the relationship between true positives and false positives for DASH across all samples in green and all samples with a tumor purity > 20% in blue. **(F)** A histogram showing the distribution of tumor purity values for alleles that were predicted incorrectly by DASH.

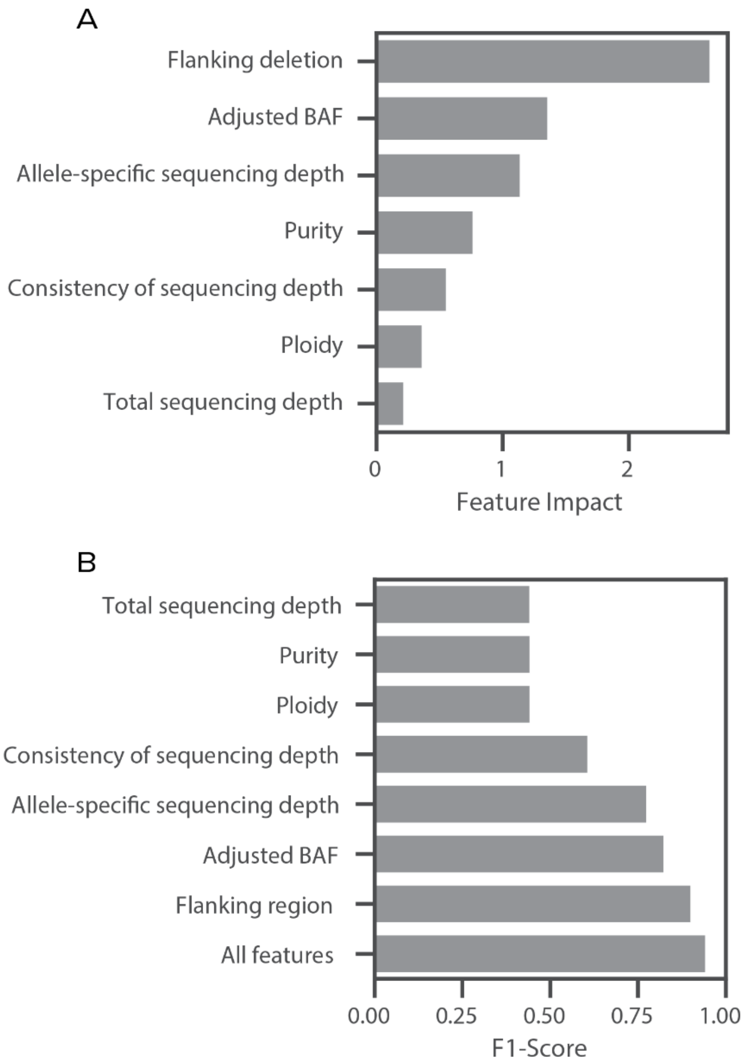

**Supplementary Figure 4. DASH feature impact. (A)** Bar plot quantifying the impact an individual feature has on the DASH algorithm. Higher feature impact denotes a feature having a higher weight in the algorithm. **(B)** Bar plot calculating the F1-score of DASH trained with only the corresponding feature.

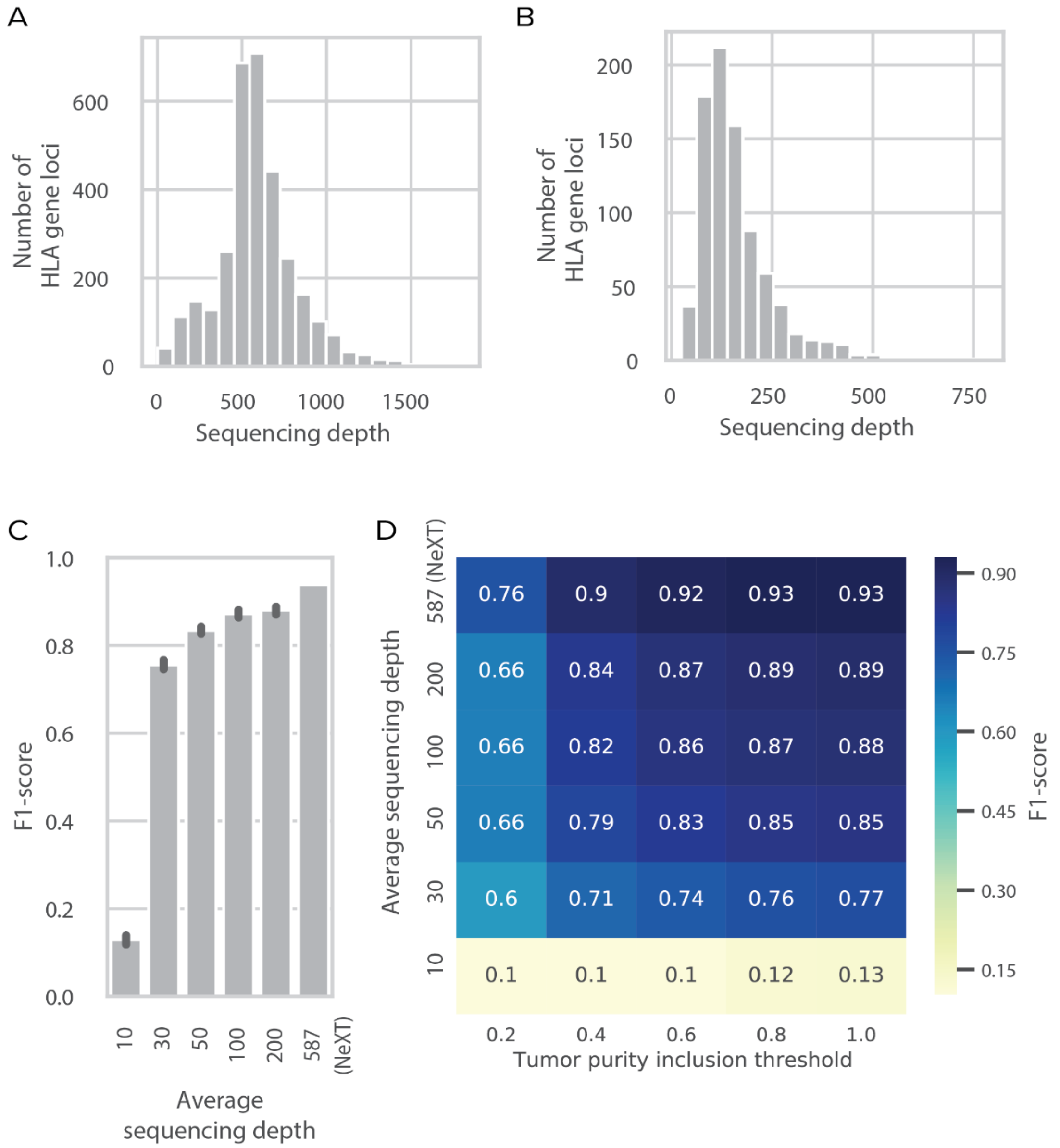


**Supplemental Figure 5. Performance of DASH with reduced sequencing depth. (A-B)** Histograms displaying the average sequencing depth found in the HLA gene loci in **(A)** ImmunoID NeXT and **(B)** standard exome sequencing. **(C)** Bar plot of the average F1-scores when DASH was run on the same set of samples across different sequencing depths. The average sequencing depth corresponding to 587 represents the average sequencing depth found in

ImmunoID NeXT samples. **(D)** Heatmap of the average F1-scores when DASH was run on the same set of samples across different sequencing depths and split by the sample's tumor purity inclusion threshold.

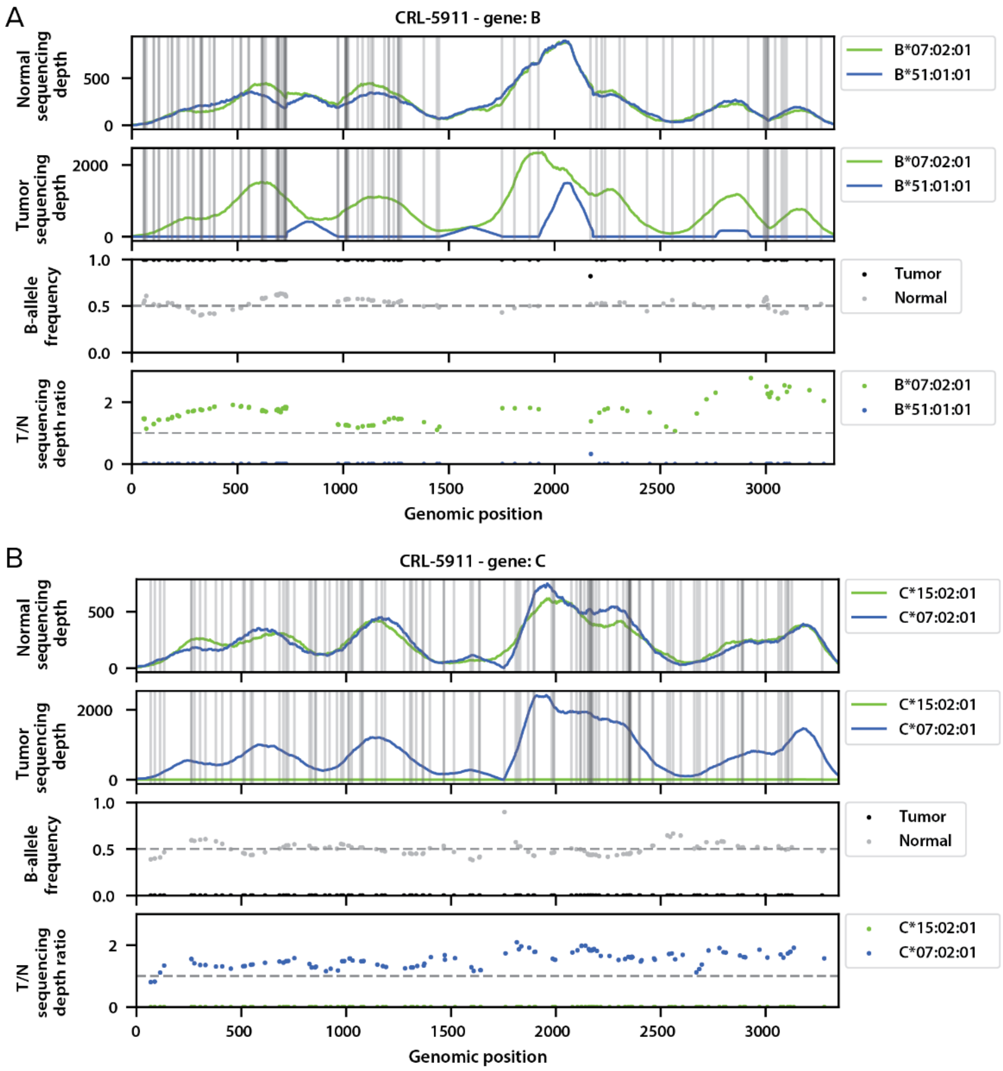


**Supplemental Figure 6. Visualization of the CRL-5911 cell-line with HLA LOH. (A-B)** Overview of features that show HLA LOH in **(A)** HLA-B and **(B)** HLA-C of the CRL-5911 cell-line.

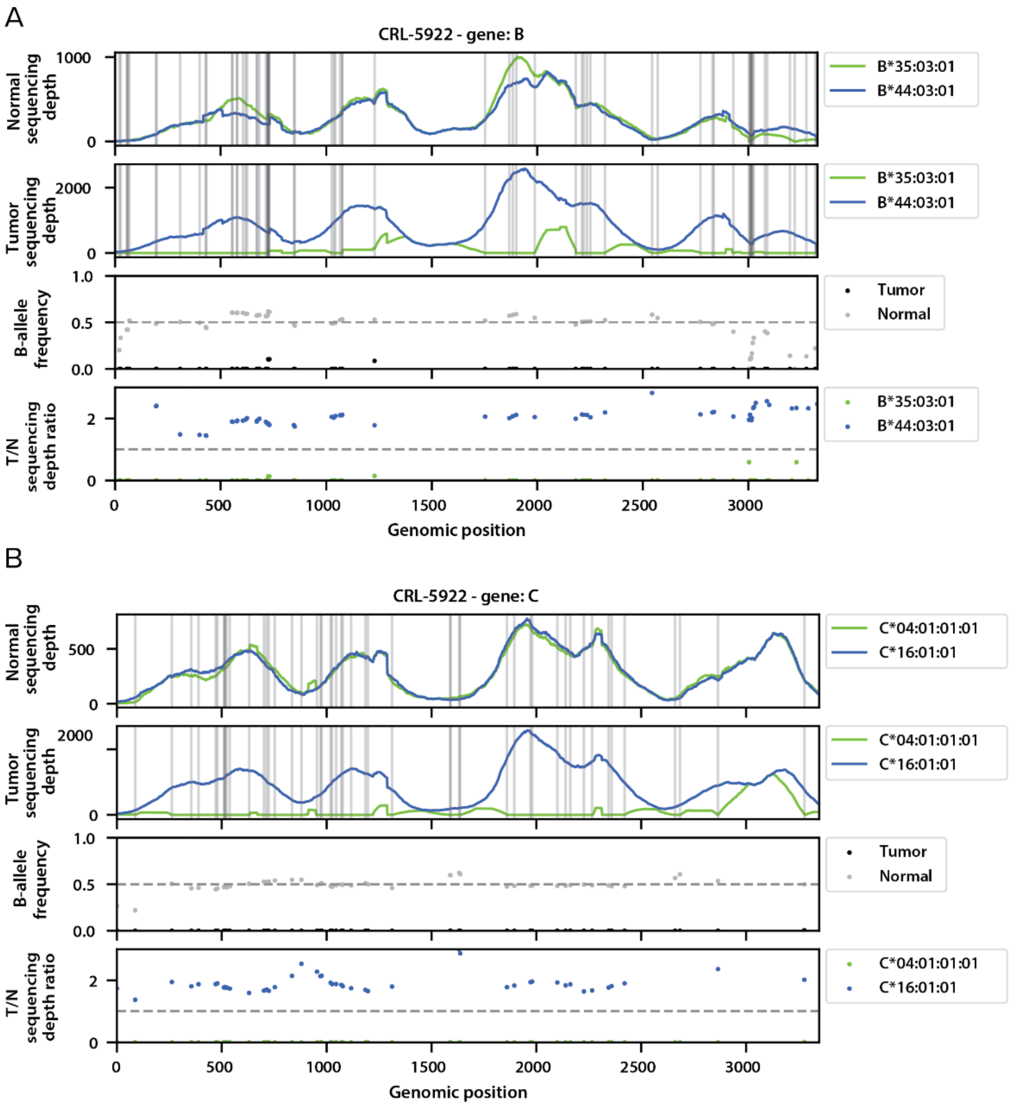


**Supplemental Figure 7. Visualization of the CRL-5922 cell-line with HLA LOH. (A-B)** Overview of features that show HLA LOH in **(A)** HLA-B and **(B)** HLA-C of the CRL-5922 cell-line.

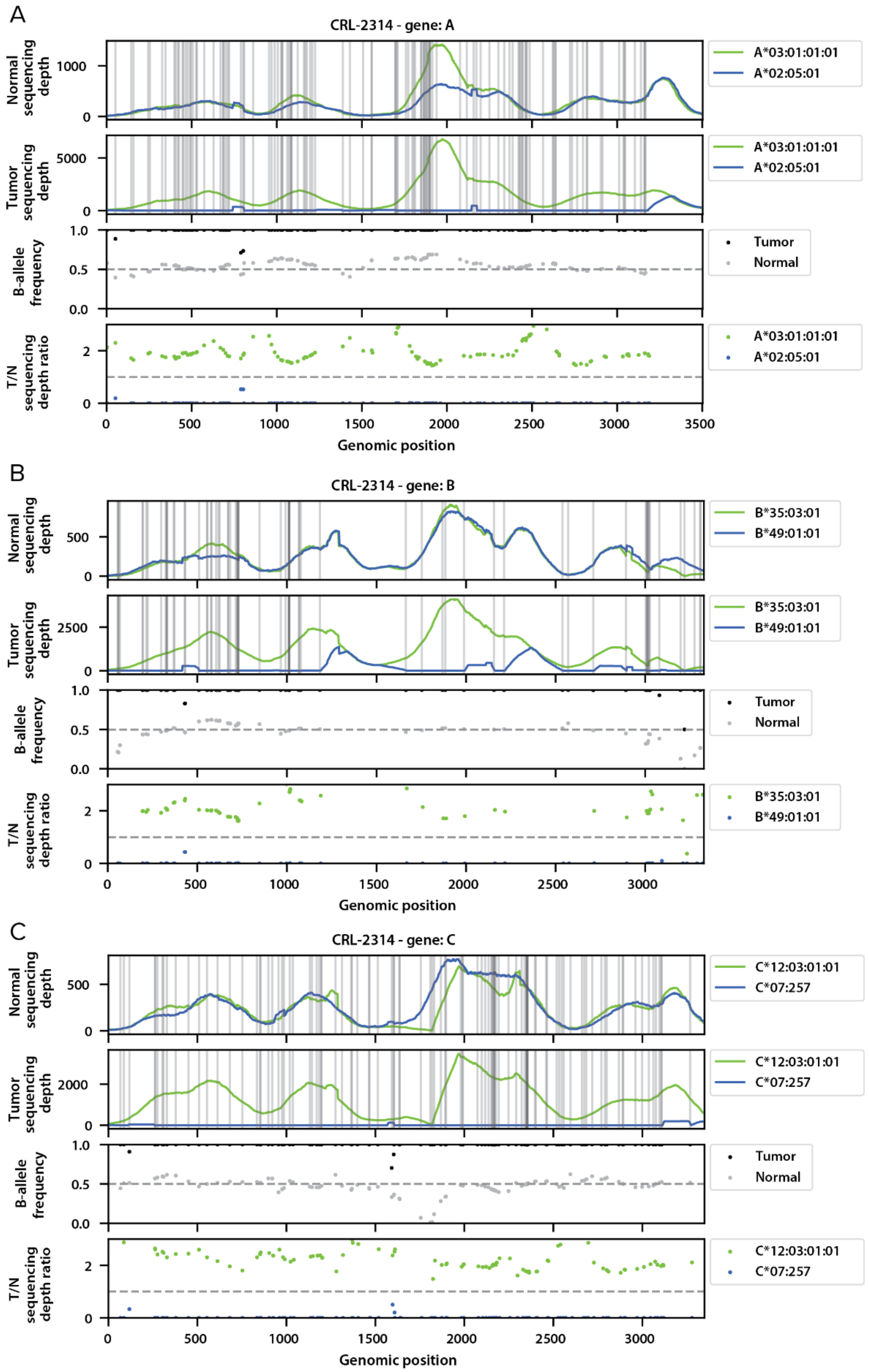


**Supplemental Figure 8. Visualization of the CRL-2314 cell-line with HLA LOH. (A-B)** Overview of features that show HLA LOH in **(A)** HLA-A, **(B)** HLA-B, and **(C)** HLA-C of the CRL-2314 cell-line.

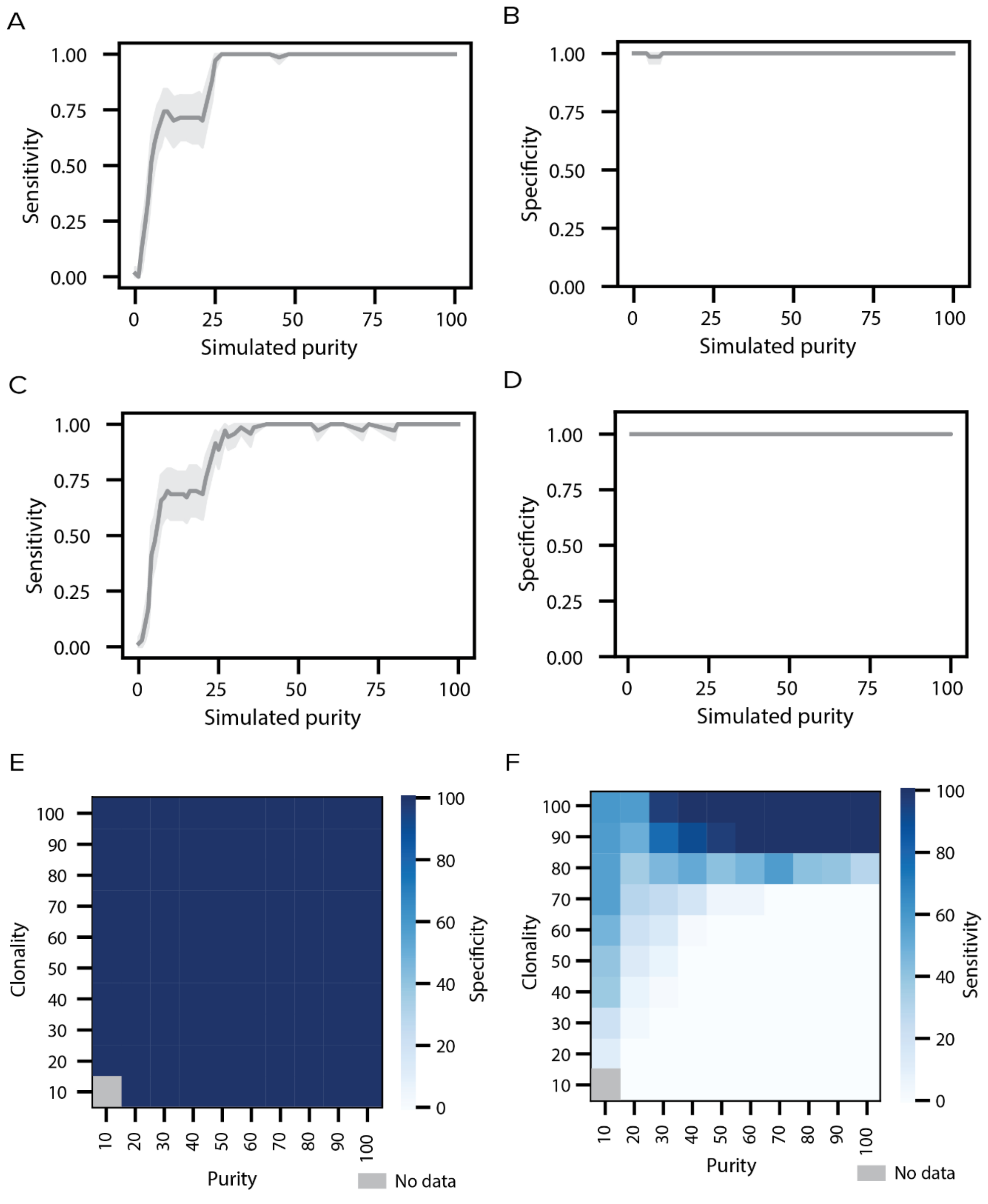


**Supplemental Figure 9. Combined LOHHLA performance on three cell-lines.**(**A-B**) Line plots showing the (**A**) sensitivity and (**B**) specificity of DASH and the (**C**) sensitivity and (**D**) specificity of LOHHLA at various purity levels with fully clonal tumors. Shaded region denotes 95% confidence. (**E-F**) Heatmaps showing the (**E**) specificity and (**F**) sensitivity of LOHHLA to capture HLA LOH in simulated samples of differing purity and clonality. Dark blue denotes high sensitivity or specificity, light blue denotes low sensitivity or specificity and grey denotes no data.

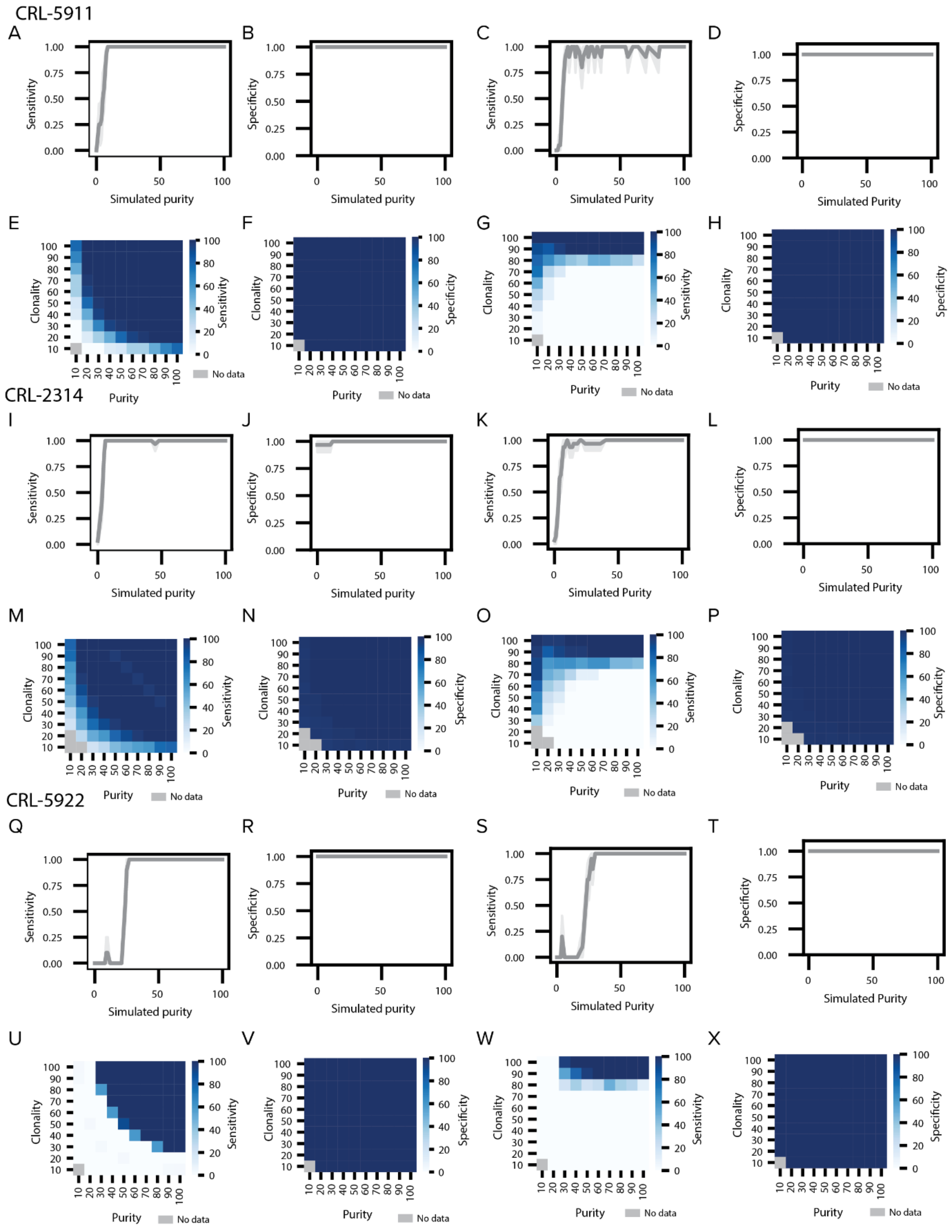


**Supplemental Figure 10. Individual DASH and LOHHLA performance on three cell-lines. (A-B, I-J, Q-R)** Line plots showing **(A, I, Q)** sensitivity and **(B, J, R)** specificity at various purity levels with fully clonal tumors using DASH. **(C-D,**

**K-L, S-T)** Line plots showing **(C, K, S)** sensitivity and **(D, L, T)** specificity at various purity levels with fully clonal tumors using LOHHLA. **(E-F, M-N, U-V)** Heatmaps displaying DASH **(E, M, N)** sensitivity and **(F, N, V)** specificity to capture HLA LOH in simulated samples of differing purity and clonality for each cell line. **(G-H, O-P, W-X)** Heatmaps displaying LOHHLA **(G, O, W)** sensitivity and **(H, P, X)** specificity to capture HLA LOH in simulated samples of differing purity and clonality for each cell line. Dark blue denotes high sensitivity or specificity, light blue denotes low sensitivity or specificity and grey denotes no data.

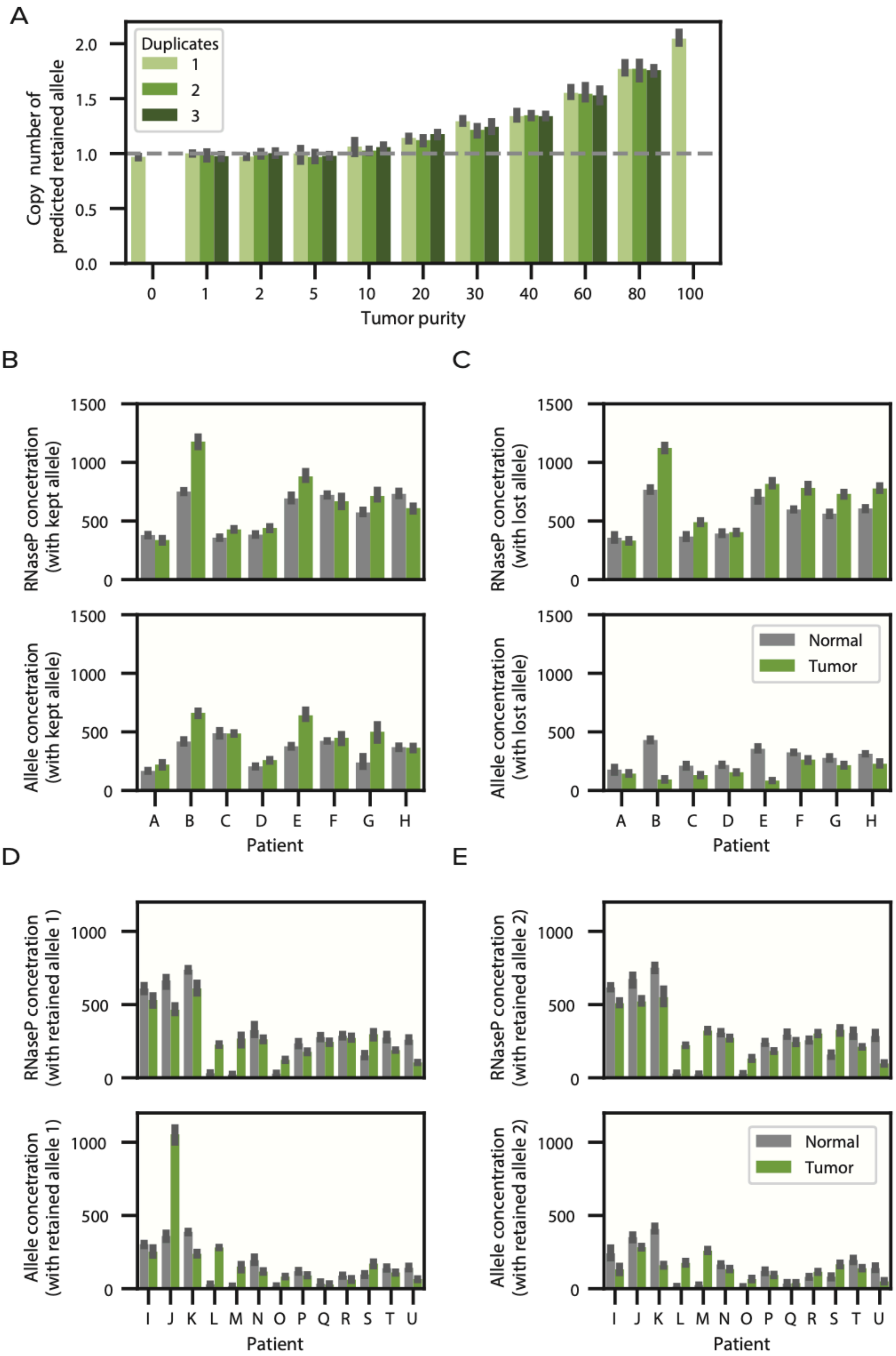
A
Duplicates
1
2
3
Copy number of predicted retained allele
2.0
1.5
1.0
0.5
0.0
0
1
2
5
10
20
30
40
60
80
100
Tumor purity
B
C
RNaseP concetration (with kept allele)
Allele concetration (with kept allele)
RNaseP concetration (with lost allele)
Allele concentration (with lost allele)
1500
1000
500
0
Normal
Tumor
A B C D E F G H
Patient
D
E
RNaseP concetration (with retained allele 1)
Allele concetration (with retained allele 1)
RNaseP concetration (with retained allele 2)
Allele concentration (with retained allele 2)
1000
500
0
I J K L M N O P Q R S T U
Patient

**Supplemental Figure 11. Allele-specific genomic validation with digital PCR. (A)** Bar plots indicating the allele-specific copy number of the predicted kept allele, relative to RNAseP, as measured by dPCR for cell line mixtures of varying tumor purities. **(B-C)** Bar plots showing RNAseP and allele concentrations for normal and tumor paired samples with HLA LOH **(B)** with the kept allele and **(C)** with the lost allele. **(D-E)** Bar plots showing RNAseP and allele concentrations for normal and tumor paired samples without HLA LOH **(D)** with the retained allele 1, **(E)** and with the retained allele 2.

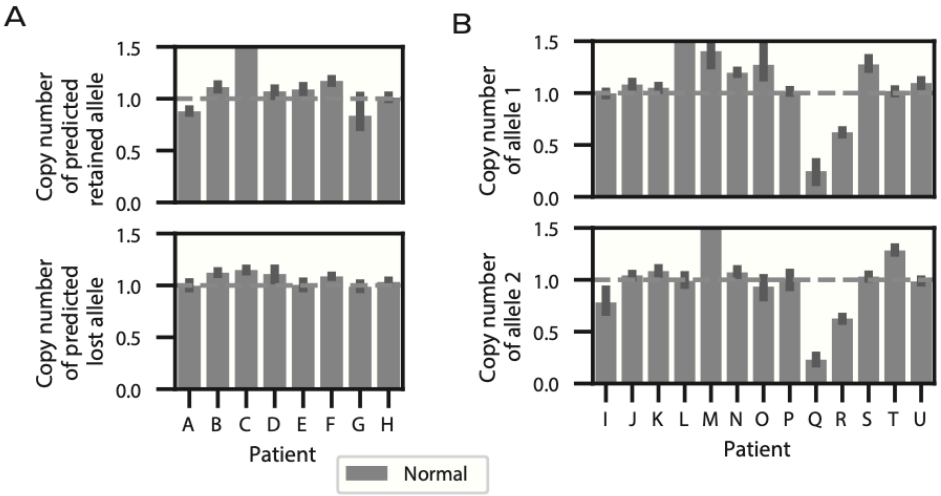


**Supplemental Figure 12. Specificities of allele-specific primers. (A)** Bar plots denoting the copy numbers of both retained and lost alleles per patient using allele-specific primers in normal diploid samples. **(B)** Bar plots denoting the copy numbers of both allele 1 and allele 2 per patient using allele-specific primers in normal diploid samples.

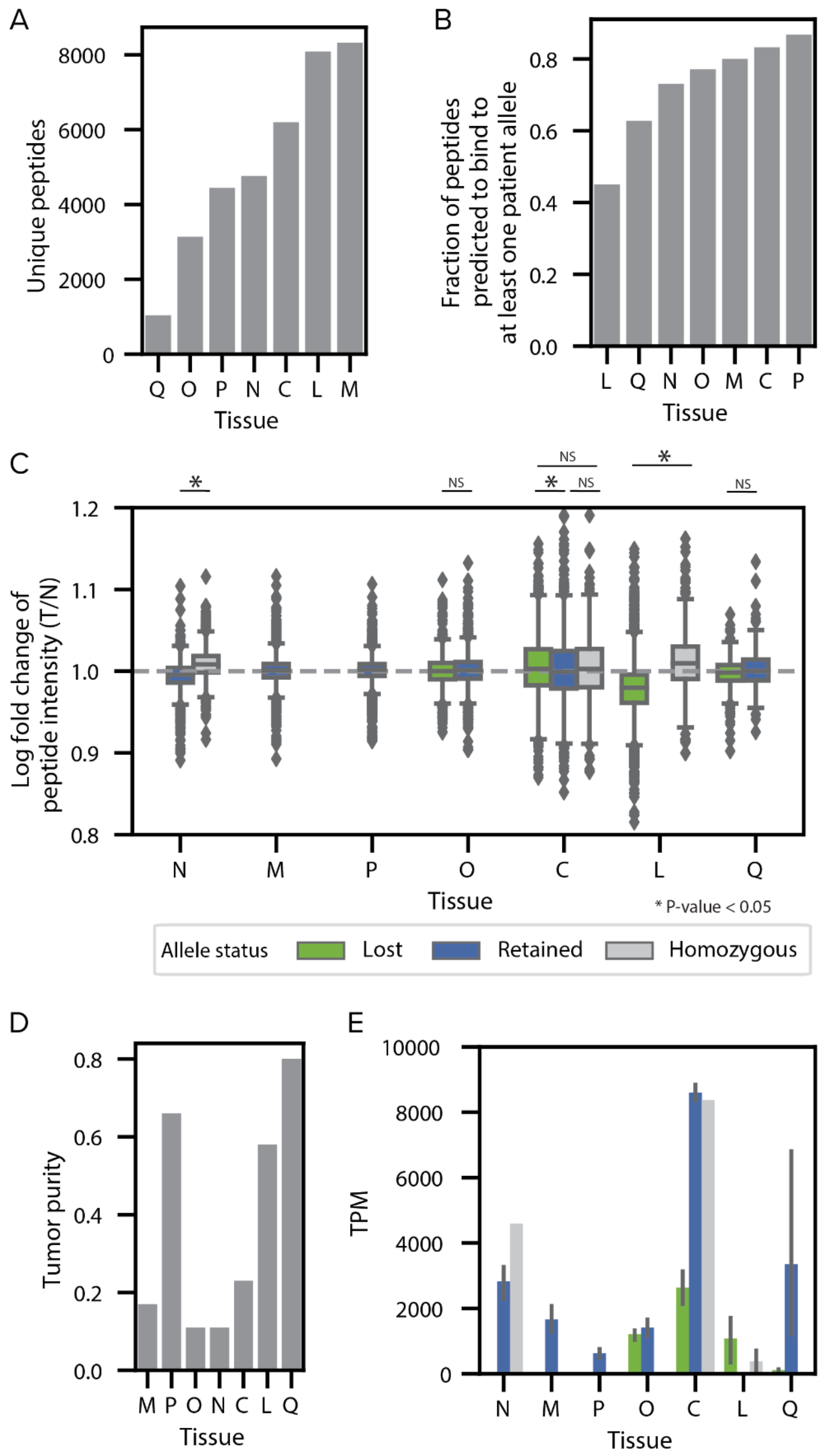


**Supplemental Figure 13. Quality and summary of quantitative immunopeptidomics data.** (**A**) A bar plot denoting the number of unique peptides identified in each quantitative immunopeptidomics experiment. (**B**) A bar plot showing the fraction of peptides in each experiment that are predicted to bind to at least one of the patient's alleles.

(**C**) Boxplots showing the distribution of log fold peptide intensities in lost, retained and homozygous alleles for each of the seven patient tissue samples. Statistical significance is assessed with a two-sided Student T-Test. (**D**) A bar plot denoting the tumor purity of each patient tumor tissue sample. (**E**) Bar plots showing the allele-specific expression distribution (TPM) of lost, retained and homozygous alleles in each patient tumor tissue sample.

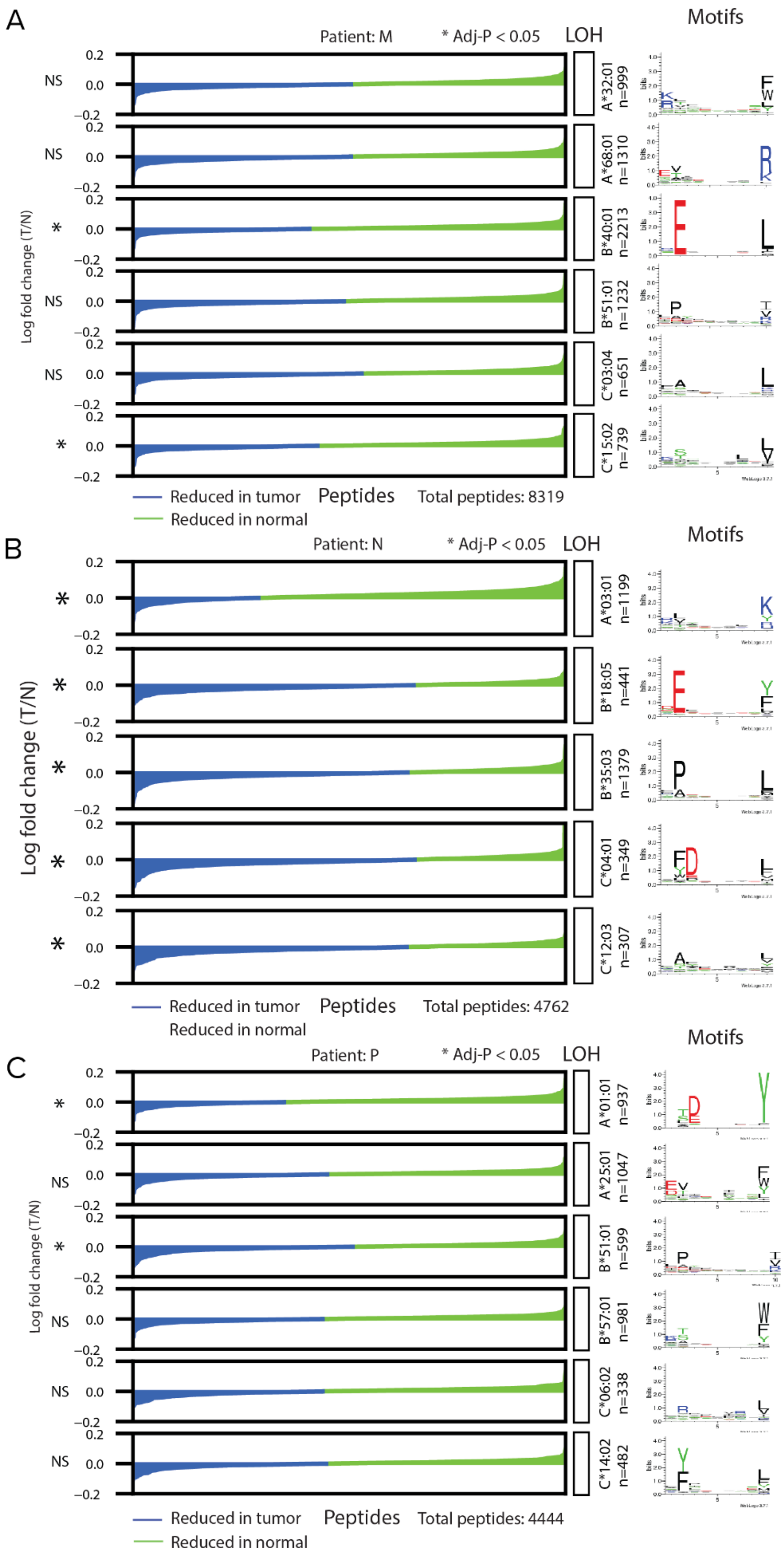
A
Patient: M
* Adj-P < 0.05
LOH
Motifs
Log fold change (T/N)
NS
NS
*
NS
NS
*
0.2
0.0
−0.2
A*32:01 n=999
A*68:01 n=1310
B*40:01 n=2213
B*51:01 n=1232
C*03:04 n=651
C*15:02 n=739
Reduced in tumor
Reduced in normal
Peptides
Total peptides: 8319
B
Patient: N
* Adj-P < 0.05
LOH
Motifs
Log fold change (T/N)
A*03:01 n=1199
B*18:05 n=441
B*35:03 n=1379
C*04:01 n=349
C*12:03 n=307
Reduced in tumor
Reduced in normal
Peptides
Total peptides: 4762
C
Patient: P
* Adj-P < 0.05
LOH
Motifs
Log fold change (T/N)
A*01:01 n=937
A*25:01 n=1047
B*51:01 n=599
B*57:01 n=981
C*06:02 n=338
C*14:02 n=482
Reduced in tumor
Reduced in normal
Peptides
Total peptides: 4444

**Supplemental Figure 14. Quantitative immunopeptidomics on control samples without predicted HLA LOH.** (**A-B**) Waterfall plots showing the log2 fold change from a normal sample to a tumor sample for peptides binding to each of the alleles in a particular patient. Blue denotes peptides that are less frequent in the tumor while green denotes peptides that are more frequent in the tumor. Grey boxes denote deleted alleles while white boxes denote retained alleles. The peptides for each allele are visualized as a motif. Statistical significance assessed using a Wilcoxon paired rank sum test. Samples shown are: (**A**) M, (**B**) N and (**C**) P.

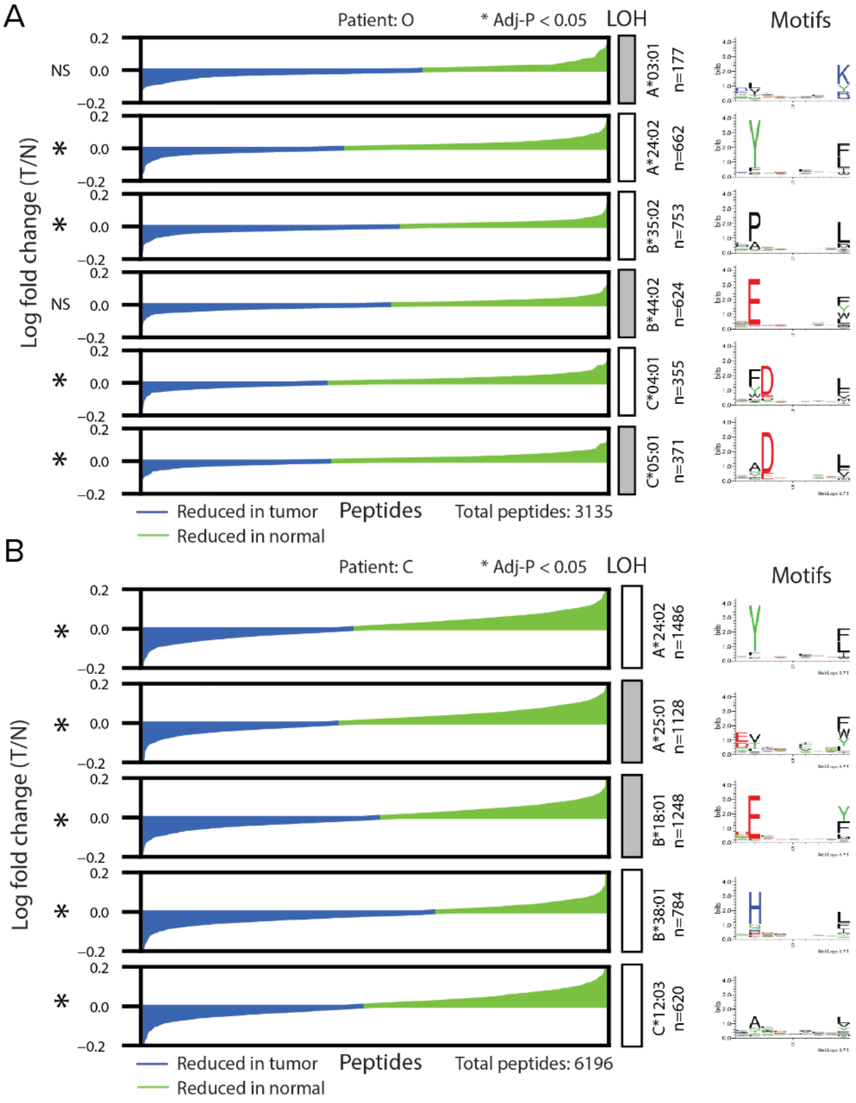


**Supplemental Figure 15. Quantitative immunopeptidomics on low tumor purity samples with predicted HLA LOH.** (**A-B**) Waterfall plots showing the log2 fold change from a normal sample to a tumor sample for peptides binding to each of the alleles in a particular patient. Blue denotes peptides that are less frequent in the tumor while green denotes peptides that are more frequent in the tumor. Grey boxes denote deleted alleles while white boxes denote retained alleles. The peptides for each allele are visualized as a motif. Statistical significance assessed using a Wilcoxon paired rank sum test. Samples shown are: (**A**) O and (**B**) C.

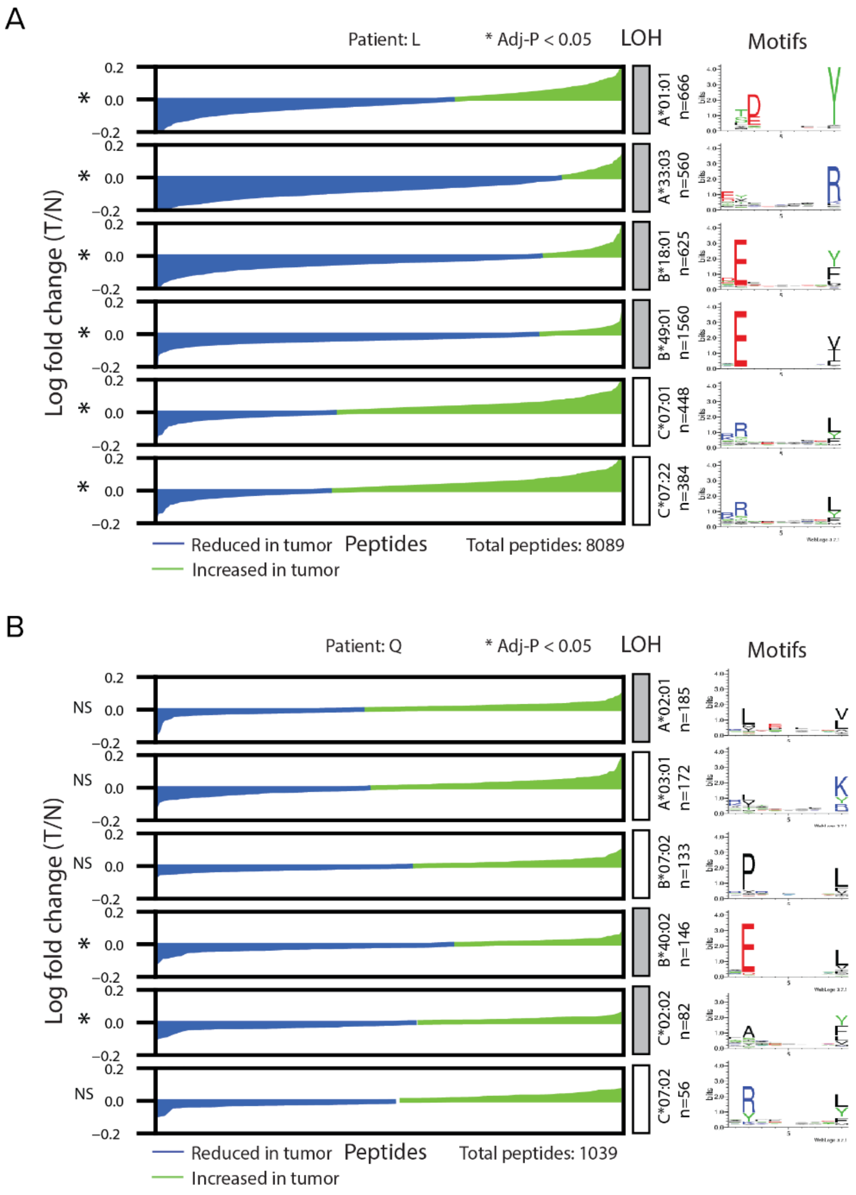


**Supplemental Figure 16. Quantitative immunopeptidomics on high tumor purity samples with predicted HLA LOH.** (**A-B**) Waterfall plots showing the log2 fold change from a normal sample to a tumor sample for peptides binding to each of the alleles in a particular patient. Blue denotes peptides that are less frequent in the tumor while green denotes peptides that are more frequent in the tumor. Grey boxes denote deleted alleles while white boxes denote retained alleles. The peptides for each allele are visualized as a motif. Statistical significance assessed using a Wilcoxon paired rank sum test. Samples shown are: (**A**) L and (**B**) Q.

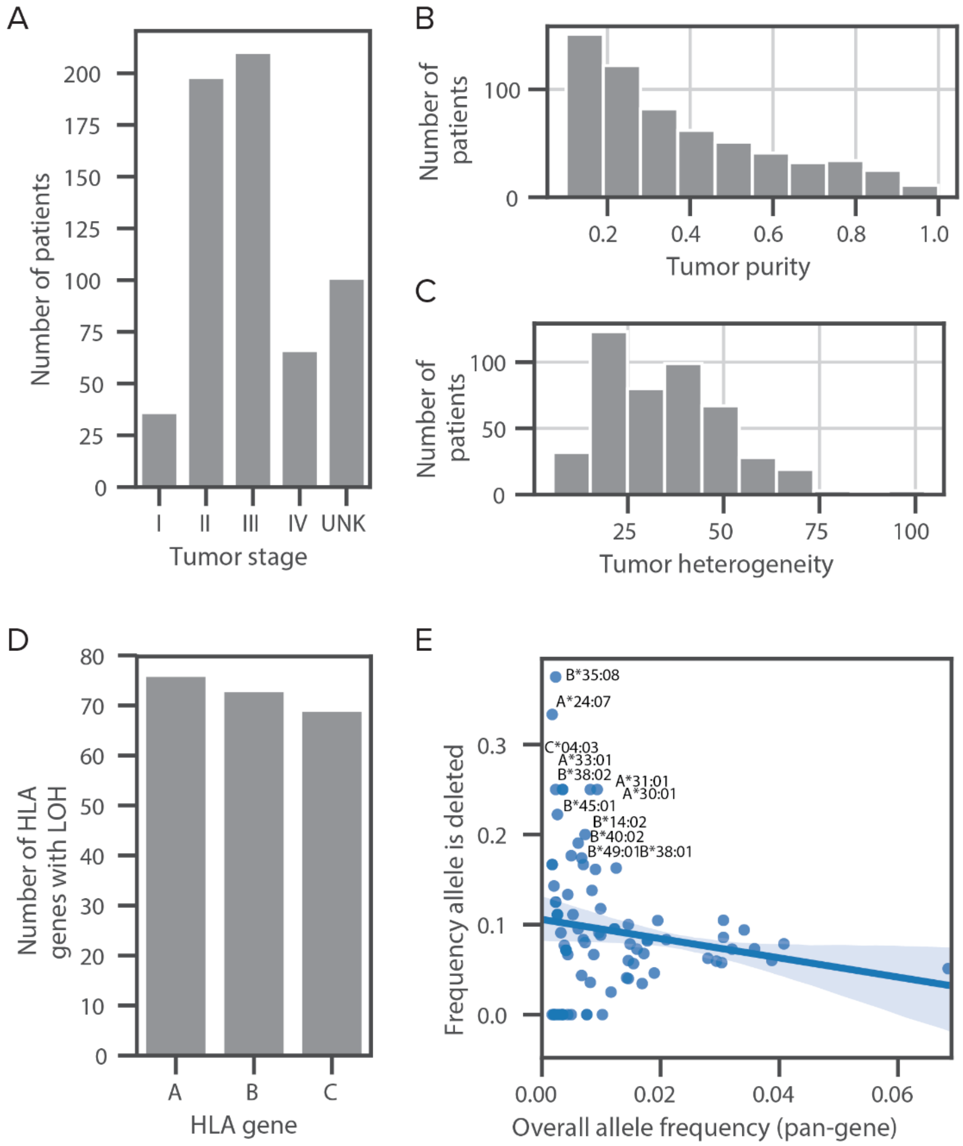


**Supplemental Figure 17. Tumor and allele breakdown across the patient cohort. (A-C)** Histograms showing the number of patients broken down by **(A)** tumor stage, **(B)** tumor purity, and **(C)** tumor heterogeneity (MATH score). **(D)** Bar plot detailing the number of HLA LOH events present in the patient cohort broken down by gene. **(E)** Scatter plot showing the frequency that an allele is deleted as a function of the overall frequency that an allele appears across the cohort.

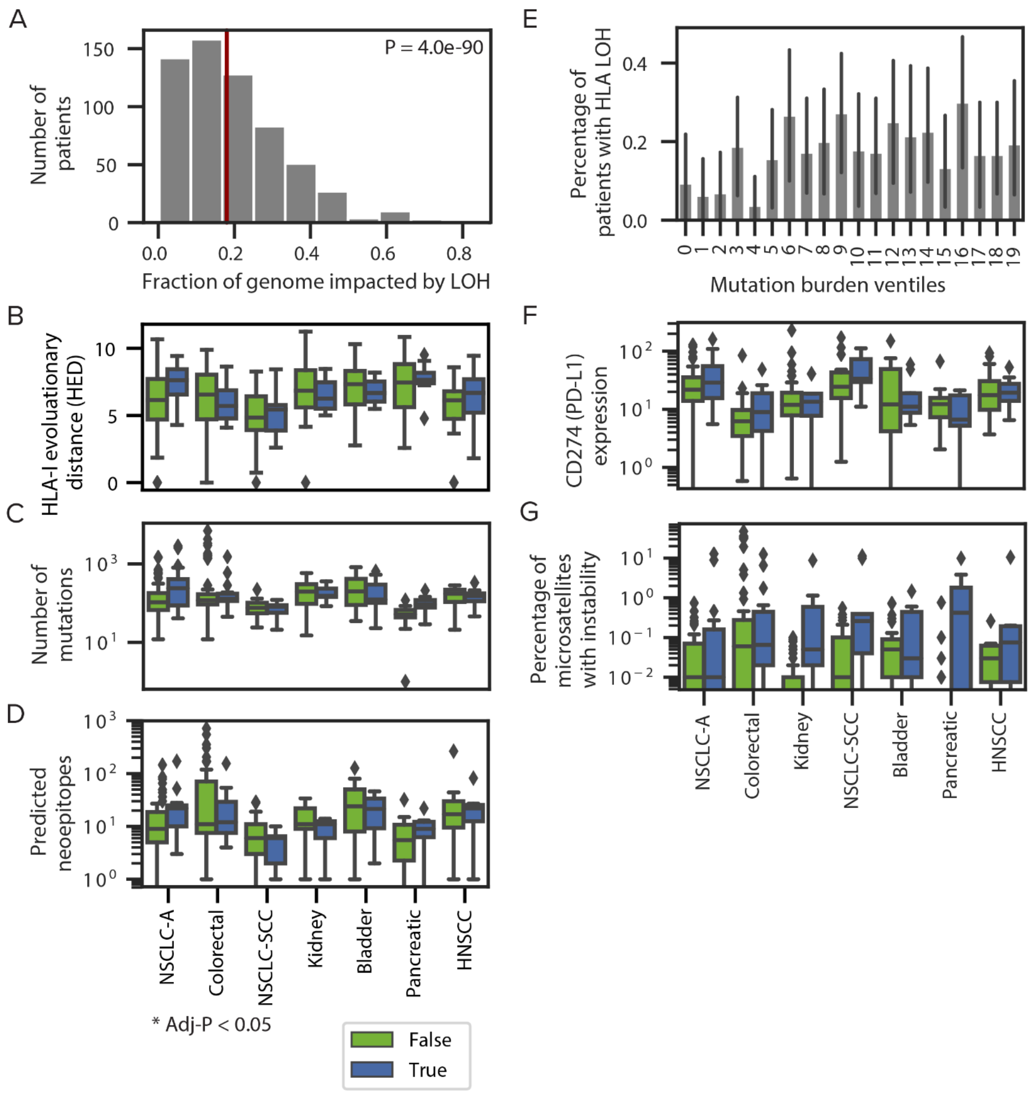


**Supplemental Figure 18. Relationship between HLA LOH and other tumor covariates.** (**A**) Histogram of the fractions of genome-wide LOH across patients in the cohort. The red-line denotes the percentage of HLA LOH in the cohort. (**B-D**) Boxplots showing the distribution of (**B**) HLA-I Evolutionary Distance (HED), (**C**) number of mutations (SNV, indel and fusion) for patients with and without HLA LOH and (**D**) predicted neoepitopes. (**E**) The percentage of patients with HLA LOH in each ventile of mutation burden pan-cancer. (**F-G**) Boxplots showing the distribution of (**F**) CD274 (PD-L1) expression and (**G**) the percentage of microsatellites with instability for patients with and without HLA LOH. For **B/C/D/F/G,** only tumor types with at least 8 patients impacted by HLA LOH are shown. Statistical analyses are performed with Mann Whitney U tests and are Bonferroni corrected.
